# Genome-wide identification of metabolic and regulatory determinants of intracellular growth in *Brucella neotomae*

**DOI:** 10.64898/2026.04.05.716554

**Authors:** Yoon-Suk Kang, James E. Kirby

**Author notes:** Corresponding author: James E Kirby.

## Abstract

We used transposon sequencing (Tn-seq) to define genetic requirements for intracellular survival of *Brucella neotomae*, a rodent-associated species. A near-saturating mutant library was subjected to selection during infection of J774A.1 macrophages, identifying 54 genes required for intracellular fitness. These included core components of the VirB type IV secretion system, multiple regulatory factors, an aquaporin gene with a strong fitness defect, and a set of metabolic genes involved in amino acid biosynthesis. Targeted mutagenesis revealed that methionine and histidine biosynthesis are indispensable for intracellular growth, whereas tryptophan biosynthesis was required for full intracellular fitness, with mutants exhibiting significant but incomplete attenuation. Notably, these auxotrophs grew normally in minimal medium under axenic conditions, indicating that their requirement is specific to the intracellular environment. Amino acid supplementation rescued intracellular growth in a concentration- and time-dependent manner, consistent with increased metabolic demand during intracellular replication. Disruption of the aquaporin gene similarly impaired intracellular survival, suggesting a role for water homeostasis during adaptation to the macrophage vacuolar environment. Beyond metabolic and osmotic adaptation, we identify OmpR1 as an upstream regulator of *B. neotomae* virulence. Biochemical, genetic, and transcriptional analyses establish a hierarchical regulatory cascade in which OmpR1 activates the BvrR/BvrS system, which in turn controls VjbR and downstream VirB expression. Under the conditions examined, OmpR1 is required for activation of this cascade. Consistent with this, OmpR1 loss is not rescued by VjbR and requires BvrR activity for restoration of intracellular growth. Phylogenetic analysis places OmpR1 in a distinct lineage relative to other well-characterized *Brucella* transcriptional regulators, suggesting that this regulatory pathway has been underappreciated across the genus. Together, these findings reveal that intracellular fitness in *Brucella* depends on metabolic capacity, osmotic homeostasis, and a hierarchical regulatory cascade centered on OmpR1.

**Author Summary:** *Brucella* species are bacteria that survive and replicate inside immune cells called macrophages, where they cause persistent infection. To live within these cells, the bacteria must carefully balance their metabolism with the expression of genes required for virulence. We used a genome-wide genetic approach to determine which genes are specifically required for intracellular survival of *Brucella neotomae*, a rodent-associated species. We found that several amino acid biosynthesis pathways, including those required to produce methionine and histidine, are essential for replication inside macrophages but are not required during growth in laboratory media. This indicates that the intracellular environment imposes nutrient limitations not apparent in culture. We also discovered that a gene encoding an aquaporin, which regulates water movement across the bacterial membrane, is critical for intracellular survival, highlighting the importance of maintaining water balance within the host cell vacuole. In addition, we identify OmpR1 as an upstream regulator that controls a hierarchical virulence cascade required for intracellular growth. Our findings show that successful infection depends on metabolic capacity, virulence regulation and water homeostasis, and provide new insight into how *Brucella* adapts to its host environment.

## Introduction

Species of the genus *Brucella* are facultative intracellular pathogens that survive and replicate within host macrophages, where they encounter a complex and dynamic environment characterized by nutrient limitation, host immune pressure (including oxidative and antimicrobial stress), and membrane stress. Successful intracellular persistence requires not only the capacity to synthesize essential metabolites, but also the ability to maintain envelope integrity and water homeostasis while sensing and adapting to conditions within the host cell through regulatory responses. While numerous studies have examined individual virulence determinants and metabolic pathways in *Brucella*, a systems-level understanding of the genetic requirements that collectively support intracellular fitness remains incomplete [1-3].

A recurring theme in *Brucella* pathogenesis is the importance of central metabolism during intracellular growth. Mutations affecting gluconeogenesis and amino acid biosynthesis have been associated with attenuation in cellular and animal infection models [4-7]. More recently, genome-wide fitness profiling approaches have reinforced this concept. Tn-seq analysis in macrophage models identified histidine biosynthesis genes as candidate intracellular fitness determinants, although without detailed validation of intracellular-specific growth requirements [8]. Similarly, organ-specific Tn-seq studies in *B. melitensis* demonstrated that tryptophan biosynthesis is required during early lung infection and that histidine biosynthetic genes contribute to in vivo fitness, underscoring the tissue-dependent nature of metabolic constraints during infection [9]. Despite these observations, it remains unclear whether such defects reflect absolute nutrient deprivation, increased metabolic demand during replication, or failures in regulatory adaptation to the intracellular environment.

In addition to metabolic enzymes, transcriptional regulators and two-component signaling systems play critical roles in enabling *Brucella* to adapt to intracellular conditions. Systems such as BvrR/BvrS and VjbR, as well as regulators linked to the stringent response and envelope stress, have been implicated in virulence and intracellular survival [6, 10-13].

Membrane adaptation may represent an additional, underappreciated component of intracellular fitness. Intracellular vacuoles undergo progressive maturation, acidification, and remodeling, processes that likely impose osmotic and envelope stress on resident bacteria [14, 15]. Although aquaporins are classically associated with water transport and osmotic regulation in bacteria [16, 17], their contribution to intracellular survival in *Brucella* has not been systematically evaluated.

Transposon sequencing (Tn-seq) provides a powerful, unbiased approach to defining genetic requirements for fitness under specific conditions, enabling the identification of metabolic functions and regulatory pathways that contribute to survival in a given environment [18, 19]. Applied to intracellular infection models, Tn-seq can reveal not only genes required for basal growth, but also those necessary for adaptation to host-specific stresses. Importantly, such analyses can distinguish between genes whose disruption causes general growth defects and those whose loss specifically impairs intracellular survival.

We have previously established *Brucella neotomae* as a genetically tractable and biologically relevant model for studying *Brucella* intracellular physiology and host-pathogen interactions. Although historically considered non-zoonotic, *B. neotomae* shares extensive genetic and functional similarity with classical pathogenic *Brucella* species and exhibits robust intracellular growth in macrophages and in vivo models. Prior work from our group has demonstrated the utility of *B. neotomae* for dissecting virulence-associated pathways, regulatory mechanisms, and host interactions, while offering experimental advantages for high-throughput genetic studies [20-23].

In this study, we employed Tn-seq to systematically identify genetic requirements for intracellular fitness of *B. neotomae* during macrophage infection. Our analysis revealed convergence of factors involved in amino acid biosynthesis, aquaporin-mediated water homeostasis, and hierarchical regulatory control of virulence gene expression.

## Results

### Genome-wide Tn-seq identifies genes required for intracellular survival of *Brucella neotomae*

To identify genetic determinants required for intracellular survival of *Brucella neotomae* in J774A.1 macrophages, we employed a genome-wide transposon sequencing (Tn-seq) approach. A saturated transposon mutant library was constructed using the Himar1 transposase system. A nourseothricin resistance cassette (NAT; nourseothricin acetyltransferase) was cloned between the Himar1 inverted repeats to enable selection of insertion mutants. The inverted repeats were modified (GCCAAC to TCCGAC) to introduce an MmeI restriction site, allowing recovery of transposon-chromosome junctions for Illumina-based sequencing [24].

Using this system, we generated an input library comprising approximately 150,000 independent *B. neotomae* mutants. This library was used to infect J774A.1 macrophages, and bacteria were recovered 48 hours post-infection to generate the output pool. Genomic DNA from both input and output pools was digested with MmeI, ligated to barcoded adapters, and selectively amplified using primers annealing to the transposon inverted repeat and adapter sequences (**Table S1**). This strategy produced sequencing-ready libraries enabling high-resolution mapping of transposon insertion sites across the *B. neotomae* genome (**Fig. 1**).

**Fig. 1.**
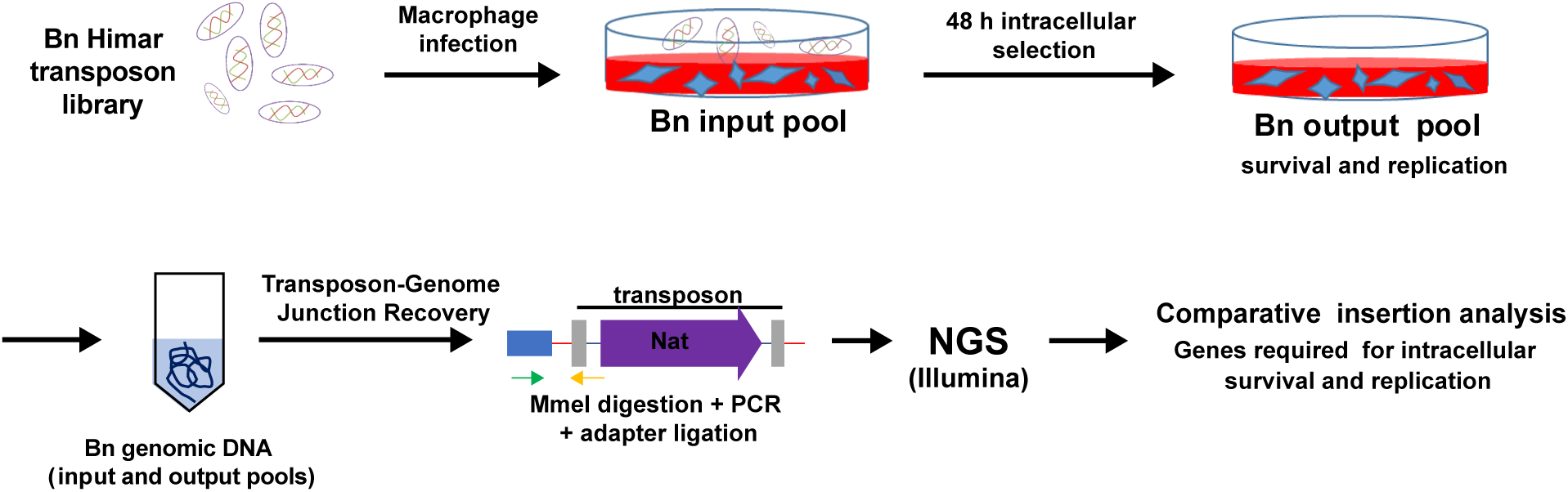
Overview of the Tn-seq workflow used to identify genes required for intracellular survival of *Brucella neotomae*. A saturated Himar1 transposon mutant library was generated and used to infect J774A.1 macrophages. Genomic DNA was isolated from the input pool prior to infection and from the output pool following 48 h of intracellular selection. Transposon-genome junctions were recovered, amplified, and subjected to high-throughput sequencing. Comparative analysis of insertion frequencies between input and output pools enabled genome-wide identification of genes required for intracellular survival and replication.

Sequencing of the input library yielded 67.5 million total reads with a mapping efficiency of 35.3%, corresponding to approximately 23.8 million informative junction reads. Of the 3,020 annotated genes, 2,787 (92.3%) contained transposon insertions, indicating near-saturating genome coverage and a well-balanced library suitable for fitness profiling. Following 48 hours of intracellular selection, sequencing depth and mapping efficiency were maintained or modestly improved (72.3 million reads; 37.8% mapping efficiency), yielding approximately 27.3 million informative reads. In contrast, the number of genes lacking insertions increased from 233 to 447, reducing genome coverage to 85.2%. Because overall sequencing depth was preserved, this loss of insertions reflects strong biological selection during intracellular growth. The additional 214 genes depleted after infection represent candidate loci that may be conditionally required for intracellular survival and replication.

### Global analysis of Tn-seq data identifies conditionally essential genes

Tn-seq analysis was performed across 3,169 annotated genetic features, including protein-coding genes as well as rRNA and tRNA loci. Both input and output pools were analyzed in three independent biological replicates, and insertion reads were quantified for each gene (**Table S2A**). Genes with extremely low read counts in the input library and consistently negligible insertion reads across all output replicates were inferred to be essential or near-essential for viability. These genes were enriched for core components of the translational machinery, including ribosomal RNAs, tRNAs, and ribosome-associated factors, and were excluded from subsequent fitness analyses.

For the remaining genes, mean insertion counts from the input and output pools were calculated, and statistical significance was assessed using p values and false discovery rate-corrected q values (**Table S2B**). Fold changes and log₂ fold changes (log2FC) were then calculated for each gene (**Table S2C**). These data were visualized using volcano plots to identify genes showing both statistically significant and condition-dependent depletion of insertions (**Fig. 2, Table S3**). Applying an FDR threshold of 10%, we identified 81 candidate genes depleted after intracellular selection. Of these, 54 genes exhibited a log2FC ≥ 2.5 and were designated as a high-confidence set of genes required for intracellular survival under the conditions tested, representing approximately 1.8% of the annotated genome (**Table S2C**). Functional categorization of the 54 strongly depleted genes identified 19 metabolic genes, 12 regulatory genes, 7 transport-associated genes, 5 components of the VirB type IV secretion system, 2 genes involved in cell envelope or polysaccharide biosynthesis, and 9 genes annotated as hypothetical proteins (**Table S2C**).

**Fig. 2.**
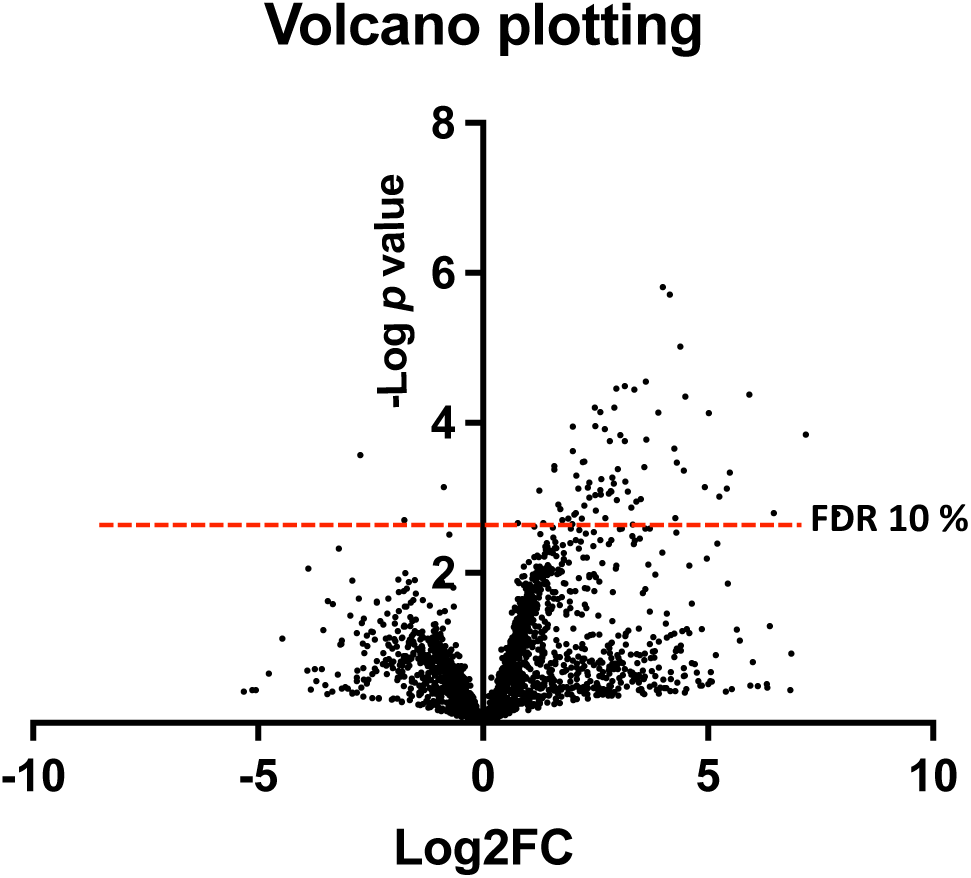
Volcano plot of Tn-seq analysis identifying genes required for intracellular survival of *Brucella neotomae*. The plot shows the log₂ fold change (Log2FC) in transposon insertion read counts between the output (intracellular) and input pools (x-axis) plotted against the −log10(p value) derived from three biological replicates per pool (y-axis). Positive Log2FC values indicate genes with reduced transposon insertions in the output pool relative to the input pool, consistent with negative selection during intracellular growth. Each point represents a single gene, quantified by aggregated insertion reads across its coding region. Statistical significance was assessed using Benjamini–Hochberg correction for multiple testing, and the dashed horizontal line denotes a false discovery rate (FDR) threshold of 10%. Genes exhibiting significant depletion of insertions in the output pool (upper right quadrant above the FDR cutoff) were classified as required for intracellular fitness, whereas genes below the threshold or near Log2FC = 0 were considered non-essential or conditionally dispensable under the experimental conditions.

To assess the biological validity of the dataset, we examined insertion profiles across the *virB* operon, which encodes the type IV secretion system and is a well-established determinant of intracellular survival. Five of the eleven *virB* genes met the criteria for significant depletion. Importantly, all *virB* genes except *virB1* showed greater than tenfold depletion of insertions following intracellular selection, with *virB7* exhibiting a particularly strong effect (log2FC = 5.2; **Table S2C**). Although some *virB* genes did not meet the final q value cutoff, the consistent depletion across the operon supports the robustness and biological relevance of the Tn-seq analysis.

### Enrichment of amino acid biosynthetic pathways among conditionally essential genes

Among the 54 genes identified as required for intracellular survival, 19 were classified as metabolism-related. Notably, pyruvate carboxylase has been reported to play a critical role in the virulence of *Brucella abortus*, and its deletion leads to attenuated infection in host cells [25]. The prominence of metabolism-associated genes in our dataset suggested that intracellular survival of *Brucella neotomae* depends not only on established virulence factors but also on the ability to meet specific anabolic demands imposed by the intracellular niche.

Further examination of these genes revealed that a substantial subset was directly or indirectly involved in amino acid biosynthesis, highlighting the importance of these pathways for intracellular survival and virulence of *Brucella*. In total, nine such genes were identified. These included two genes involved in methionine biosynthesis, three genes associated with histidine biosynthesis, and four genes belonging to the tryptophan biosynthetic pathway. Based on these findings, we next examined the contribution of these pathways to intracellular growth using targeted mutants.

### Methionine and histidine biosynthesis are required for intracellular growth

To investigate the contribution of individual amino acid biosynthetic pathways to intracellular infection by *B. neotomae*, we constructed mutants in genes involved in the biosynthesis of methionine (*metH*), histidine (*hisG*), and tryptophan (*trpD*) and used these strains to infect J774A.1 macrophages. As shown in Fig. 3A, the *metH*- and *hisG*-deficient strains exhibited complete inhibition of intracellular growth at 48 hours post-infection, comparable to the phenotype observed for the *virB4* mutant. In contrast, the *trpD*-deficient strain retained limited intracellular growth, with an approximately 96% reduction relative to the wild-type strain.

**Fig. 3.**
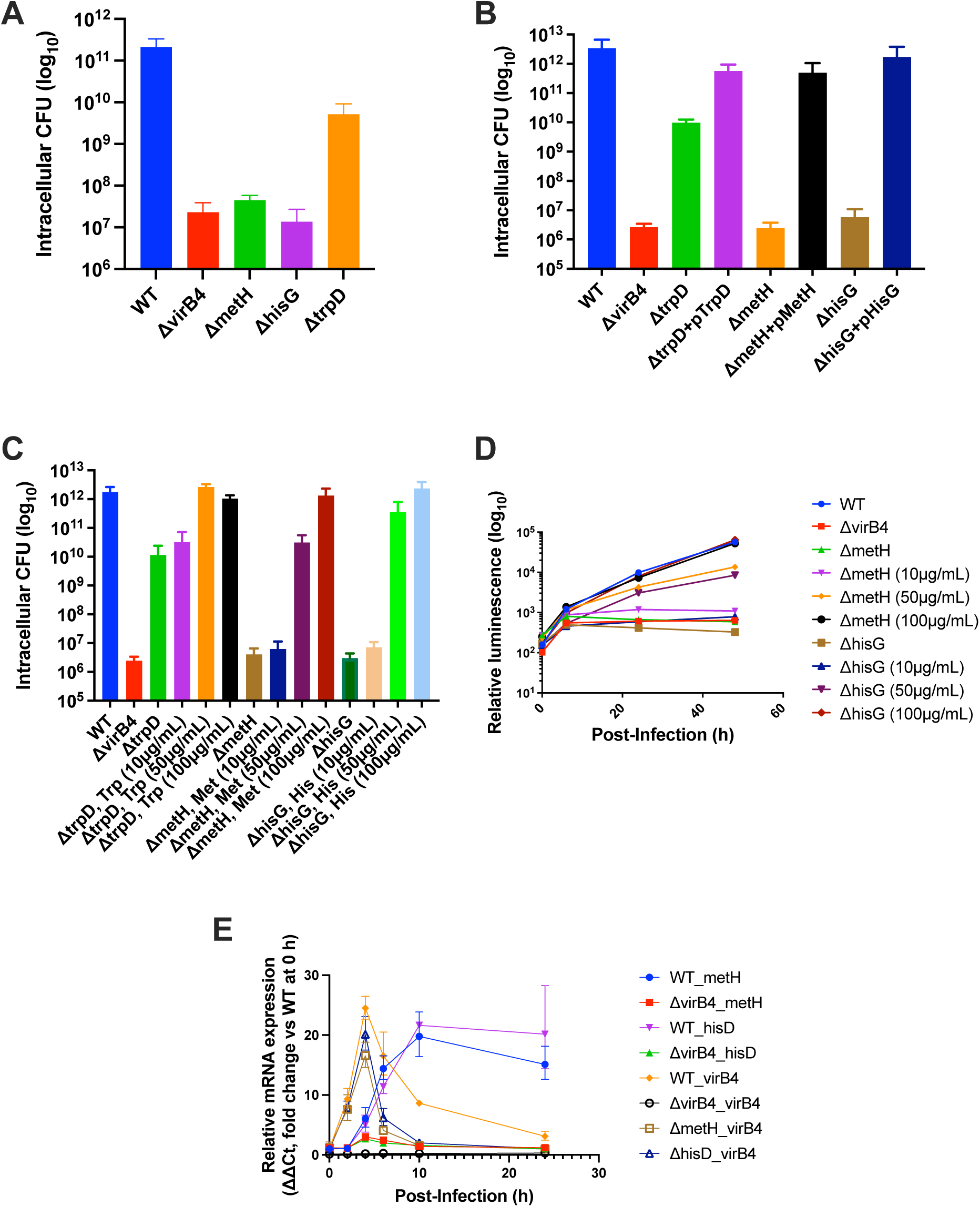
Contribution of amino acid biosynthesis genes to intracellular growth of *Brucella neotomae*. **(A)** J774A.1 macrophages were infected with WT, Δ*virB4*, Δ*metH*, Δ*hisG*, or Δ*trpD* strains at a multiplicity of infection (MOI) of 10. Intracellular CFU were quantified by CFU enumeration at 48 h post-infection. **(B)** J774A.1 macrophages were infected with WT, Δ*virB4*, Δ*metH*, Δ*hisG*, and Δ*trpD* strains and their corresponding plasmid-complemented derivatives at an MOI of 10, and intracellular CFU were determined at 48 h post-infection. **(C)** J774A.1 macrophages were infected as in (A) in the presence of methionine, histidine, or tryptophan supplementation (10, 50, or 100 µg/mL, as indicated), added at the time of infection. Intracellular CFU were enumerated at 48 h post-infection. **(D)** Intracellular growth of luminescent WT, Δ*virB4*, Δ*metH*, and Δ*hisG* strains was monitored over time following infection at an MOI of 10 by measuring relative luminescence units (RLU), which correlate with intracellular CFU as described previously. Luminescence was measured at 0, 6, 24, and 48 h post-infection. **(E)** Expression of selected amino acid biosynthesis and regulatory genes during intracellular infection was assessed by quantitative RT-PCR. J774A.1 macrophages were infected at an MOI of 100, RNA was harvested at the indicated time points post-infection, and relative mRNA levels were calculated using the ΔΔCt method. For each gene analyzed, expression in the corresponding wild-type (WT) strain at 0 h served as the reference condition. Data are plotted as fold change relative to this gene-specific WT 0 h baseline. Labels indicate the infecting strain background (WT or indicated in-frame deletion mutant), followed by the gene whose expression was quantified by RT-qPCR. Data represent the mean ± SD of n = 3 biological replicates for panels (A–D) and n = 2 biological replicates for panel (E).

To confirm that the intracellular growth defects observed in the amino acid biosynthesis mutants were attributable to the targeted deletions, complemented strains were constructed for each mutant. Intracellular growth was fully restored in these strains (**Fig. 3B**), confirming that the observed phenotypes resulted from these mutations.

### Concentration-dependent rescue by amino acid supplementation

To further assess whether intracellular growth defects resulted from amino acid limitation within host cells, macrophage infections were performed in the presence of increasing concentrations of the corresponding amino acids, 10, 50, or 100 µg/mL. Intracellular bacterial burden was quantified by CFU enumeration at 48 h post-infection. Supplementation at 10 µg/mL failed to rescue intracellular growth for any mutant strain. In contrast, tryptophan supplementation fully restored intracellular growth of the *trpD* mutant at 50 µg/mL, whereas methionine and histidine required higher concentrations, 100 µg/mL, to completely restore intracellular growth to wild-type levels (**Fig. 3C**).

Consistent with endpoint CFU measurements, time-course analysis of intracellular growth using luminescence revealed that methionine- and histidine-deficient mutants supplemented with 50 µg/mL exhibited partial rescue, whereas supplementation at 100 µg/mL supported markedly greater intracellular growth during the later stages of infection (**Fig. 3D**). These results indicate that methionine and histidine availability becomes increasingly limiting as intracellular infection progresses.

### Histidine biosynthesis is required independently of purine and pyrimidine metabolism

Because *hisG*, which catalyzes the first committed step in histidine biosynthesis, generates the intermediate AICAR that can feed into purine metabolism, we generated a Δ*hisD* mutant in which only the terminal step of histidine synthesis is disrupted. Intracellular growth of Δ*hisG* and Δ*hisD* mutants was monitored in J774A.1 macrophages over time following infection (**Fig. S5**). Intracellular growth was completely abolished in both mutants, demonstrating that the requirement for histidine biosynthesis during intracellular growth is not due to indirect effects on purine or pyrimidine metabolism. Together, these results establish that histidine biosynthesis is required for intracellular infection by *Brucella neotomae* and, in combination with methionine and tryptophan auxotrophy phenotypes, highlight a broader dependence on amino acid biosynthetic pathways during intracellular growth.

### Amino acid biosynthesis is uncoupled from VirB induction but linked to BCV maturation

To determine whether amino acid auxotrophy affected virulence gene expression, we examined the time-dependent expression of *virB4*, *metH*, and *hisD* following infection of J774A.1 macrophages (**Fig. 3E**). In the wild-type strain, *virB4* expression began to increase at 2 hours post-infection, peaked at 4 hours, and subsequently declined. Importantly, *virB4* expression in the *metH* and *hisD* mutant strains was comparable to that in the wild type strain, indicating that amino acid auxotrophy does not impair induction of the VirB virulence system. In contrast, expression of *metH* and *hisD* increased beginning at 4 hours post-infection and remained elevated at later time points. Notably, expression of these genes was markedly reduced in the *virB4* mutant strain. These results suggest that methionine and histidine biosynthesis is required during later stages of intracellular infection, rather than during early entry or VirB activation.

### Rsh-dependent regulation of intracellular metabolism and virulence

Among the metabolism-related genes identified as required for intracellular growth, the *rsh* gene, encoding a GTP pyrophosphokinase responsible for synthesis of the alarmone, ppGpp, was of particular interest. Previous studies have consistently shown that *Brucella rsh* mutant strains display pronounced defects in intracellular growth in both macrophages and mouse infection models [6, 26, 27]. In these studies, genes regulated by *rsh* were enriched for amino acid biosynthesis pathways, including threonine and histidine [27]. In addition, the *virB* operon was shown to be under *rsh* control and was not expressed in *rsh* mutant strains [6, 26]. Likewise, genes involved in methionine biosynthesis were reported to be regulated by *rsh*, and the growth defect caused by *rsh* deficiency in minimal medium was rescued by methionine supplementation [26].

Based on these observations, we generated an *rsh* mutant strain in *B. neotomae* and infected J774A.1 macrophages to assess intracellular growth. Consistent with previous studies, the *rsh* mutant exhibited complete inhibition of intracellular growth in macrophages, whereas this defect was restored in the complemented strain (**Fig. 4A**). We next monitored the time-dependent expression of *virB4* and *rsh* in the wild-type, *virB4*, and *rsh* mutant strains following infection (**Fig. 4B**). Expression of *rsh* peaked at 4 hours post-infection and remained elevated at 24 hours. In contrast, in the *virB4* mutant strain, *rsh* expression at 4 hours was reduced to approximately half of the wild-type level and was no longer detectable at 24 hours post-infection. Consistently, *virB4* expression was not detected at 4 hours post-infection in the *rsh* mutant strain. Taken together, these results indicate that, in *B. neotomae*, *rsh* influences the VirB system.

**Fig. 4.**
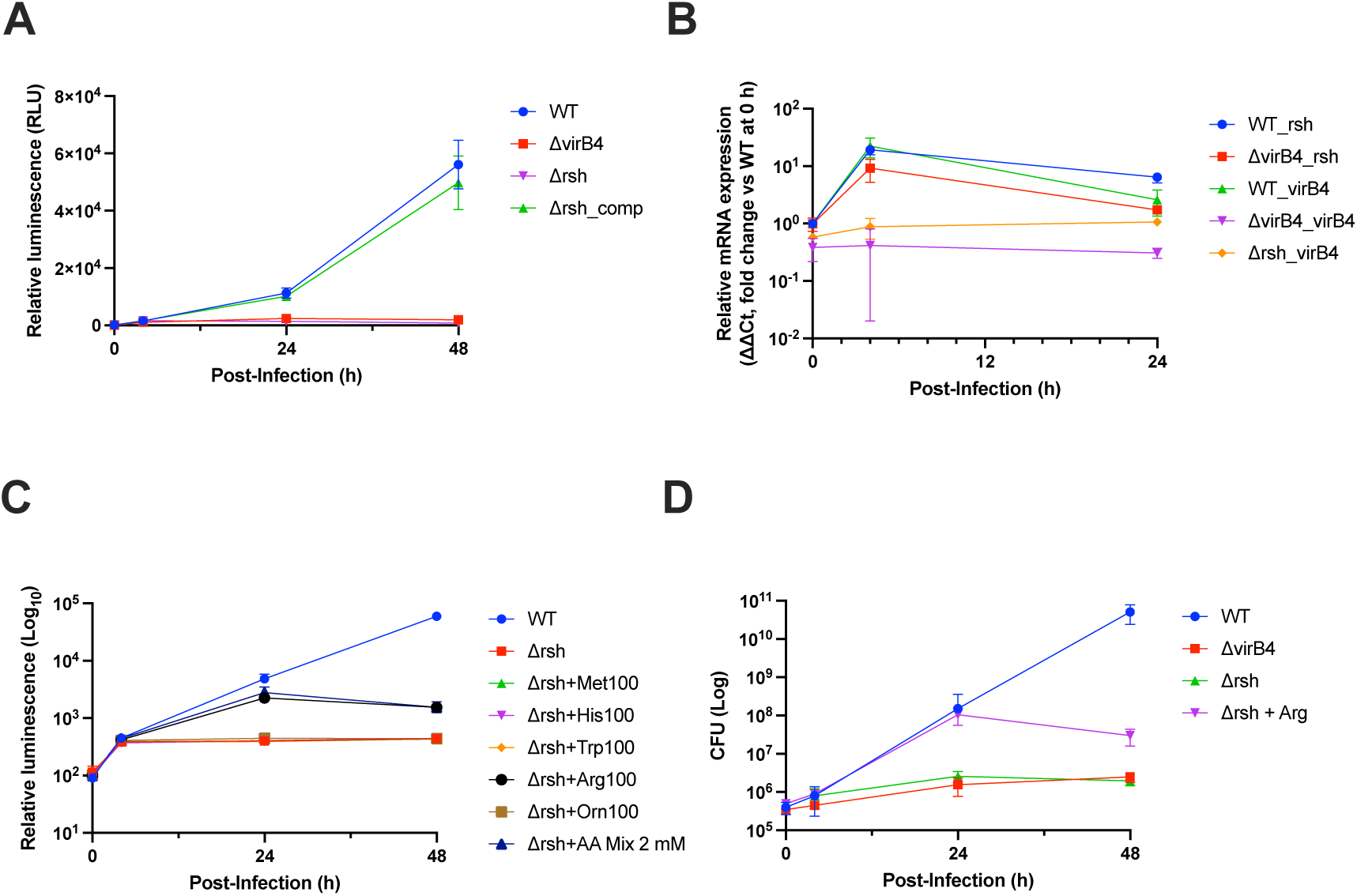
Role of the *rsh* gene in intracellular growth of *Brucella neotomae* and metabolic pathways influenced by *rsh* expression. **(A)** J774A.1 macrophages were infected with WT, Δ*virB4*, Δ*rsh*, or the Δ*rsh* complemented strain at an MOI of 10. Intracellular growth was monitored over time by measuring bacterial luminescence. Data represent the mean ± SD of n = 3 biological replicates. **(B)** To assess expression of *rsh* and *virB4* during intracellular infection, J774A.1 macrophages were infected with WT, Δ*virB4*, or Δ*rsh* strains at an MOI of 100. RNA was harvested at the indicated time points (0, 4, and 24 h post-infection), and relative mRNA levels were quantified by qRT-PCR using the ΔΔCt method. For each gene analyzed, expression in the corresponding WT strain at 0 h served as the reference condition. Data represent the mean ± SD of n = 2 biological replicates. **(C, D)** Intracellular growth of the Δ*rsh* strain was evaluated in the presence of amino acid supplementation. The Δ*rsh* strain was supplemented at the time of infection with methionine, histidine, tryptophan, arginine, or ornithine (100 µg/mL), or with a 2 mM amino acid mixture, and used to infect J774A.1 macrophages at an MOI of 10. Intracellular growth was monitored at 0, 4, 24, and 48 h post-infection by measuring luminescence (C) and CFU enumeration (D). Luminescence data represent the mean ± SD of n = 4 biological replicates, and CFU data represent the mean ± SD of n = 2 biological replicates.

We next examined whether the intracellular growth defect of the *rsh* mutant could be rescued by methionine or other amino acids (**Fig. 4C**). To this end, an amino acid mixture (2 mM) or individual amino acids (100 µg/mL) were added during infection of J774A.1 macrophages with the *rsh* mutant strain. Time-course analysis of intracellular growth showed that, at 24 hours post-infection, the *rsh* mutant supplemented with either the amino acid mixture or arginine exhibited intracellular growth levels comparable to those of the wild-type strain. However, by 48 hours post-infection, intracellular growth was no longer sustained. To confirm these observations using an independent viability readout, CFU assays were also performed and yielded consistent results (**Fig. 4D**). Taken together, these findings indicate that arginine supplementation can transiently support intracellular growth of the *rsh* mutant.

To determine whether the intracellular growth defects observed in the Δ*rsh*, Δ*metH*, and Δ*hisG* mutant strains reflect a generalized growth defect under nutrient-limiting conditions, we assessed their growth in axenic M9 minimal medium over time. Under the defined in vitro conditions used in this study, all mutant strains exhibited growth comparable to the wild-type strain by 48 hours (**Fig. S6**). These findings indicate that the intracellular growth defects are not attributable to a generalized inability to grow under nutrient-limiting conditions but instead reflect a specific impairment in adaptations required for intracellular survival within host cells.

### Genes involved in transport

Among the nine transport-associated genes depleted during intracellular selection, we selected the aquaporin gene *aqpZ* for targeted validation. Deletion of *aqpZ* produced a marked intracellular growth defect at 48 h post-infection, approaching the magnitude observed for the Δ*virB4* control, and this phenotype was restored by plasmid-based complementation (**Fig. S7**). In contrast, the Δ*aqpZ* strain exhibited no growth defect in TSB or defined M9 minimal medium. However, the mutant showed increased sensitivity to elevated NaCl, with a pronounced defect at higher salt concentrations in TSB and at lower NaCl concentrations in defined M9 medium. Taken together, these data indicate that *aqpZ* is dispensable for basal growth under standard laboratory conditions but contributes to osmotic stress resistance and intracellular growth.

### Genes involved in regulatory functions

After identifying metabolic pathways required for intracellular survival, we next examined regulatory genes identified in the Tn-seq analysis (**Table S2C**). Among these, *deoR/glpR* and *vjbR* have previously been shown to influence *Brucella* virulence [11, 28-31]. These genes encode single-component transcriptional regulators belonging to the GntR and LuxR families, respectively. *rpoN* encodes the alternative sigma factor σ⁵⁴, which has been implicated in global transcriptional regulation but was previously reported not to be required for intracellular growth in macrophage or mouse infection models [32]. OmpR1 also stood out as a predicted member of a two-component regulatory system. OmpR1 is an OmpR-family response regulator, sharing conserved receiver and DNA-binding domains with canonical OmpR proteins. Genomic organization and sequence homology indicate that OmpR1 is paired with an adjacent EnvZ-like histidine kinase, designated EnvZ1, forming a putative OmpR1/EnvZ1 two-component system.

To assess the contribution of these regulators to *B. neotomae* virulence, we generated mutant strains lacking *deoR/glpR*, *vjbR*, *ompR1*, and *rpoN*. J774A.1 macrophages were infected with each mutant strain, and intracellular survival was evaluated by CFU enumeration at 48 h post-infection (**Fig. 5A**). Deletion of *rpoN* had no detectable effect on intracellular growth. In contrast, loss of *deoR/glpR*, *vjbR*, or *ompR1* resulted in complete inhibition of intracellular growth, indicating that these regulators are required for survival and replication within host cells.

**Fig. 5.**
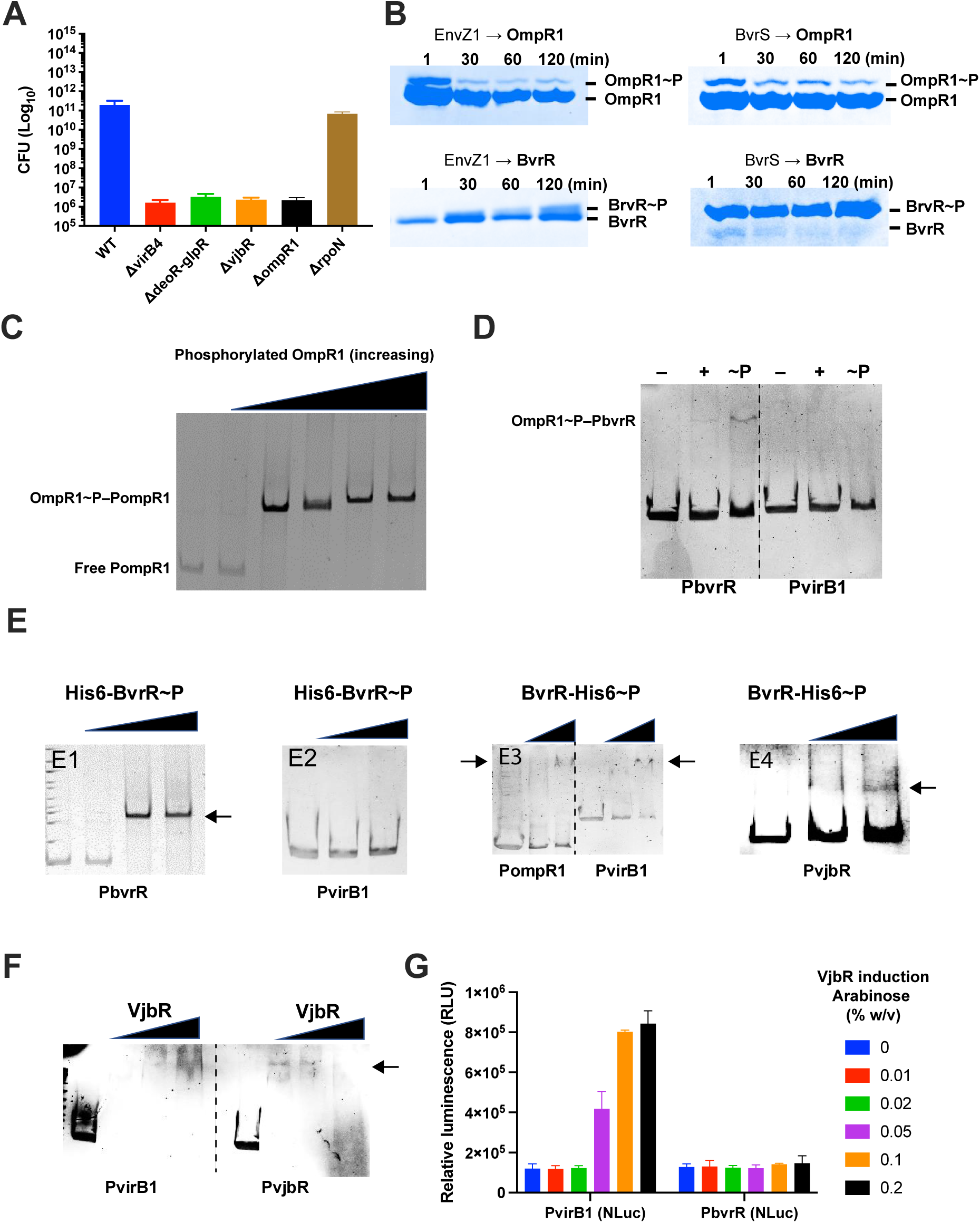
Role of essential regulatory genes in intracellular growth and virulence gene regulation in *B. neotomae*. **(A)** To determine the contribution of essential regulatory genes to intracellular survival, J774A.1 macrophages were infected at an MOI of 10 with wild-type (WT) *B. neotomae* or mutant strains lacking *virB4*, *deoR/glpR*, *vjbR*, *ompR1*, or *rpoN*. At 48 h post-infection, macrophages were lysed and intracellular bacteria were recovered for enumeration of colony-forming units (CFU) by serial dilution and plating. Data represent the mean ± SD of n = 3 biological replicates. **(B)** Phosphorelay and cross-phosphorylation between the OmpR1/EnvZ1 and BvrR/BvrS two-component systems. EnvZ1 or BvrS (0.7 µM) was autophosphorylated by addition of 50 µM ATP and then incubated with OmpR1 or BvrR (3–4 µM) for 0–120 min. Reactions were terminated by addition of sample buffer and analyzed by Phos-tag SDS–PAGE to resolve phosphorylated (OmpR1∼P, BvrR∼P) and unphosphorylated species. Lane 1, 1 min; lane 2, 30 min; lane 3, 60 min; lane 4, 120 min. Panels represent independent experiments; differences in band separation reflect gel-to-gel migration variability. In the BvrS → BvrR reaction, unphosphorylated BvrR decreased as BvrR∼P accumulated over time. **(C)** Electrophoretic mobility shift assay showing dose-dependent binding of phosphorylated OmpR1 to the *ompR1* promoter (P_ompR1), resulting in formation of a shifted OmpR1∼P–P_ompR1 complex. EnvZ1 (0.7 µM) was autophosphorylated with 50 µM ATP and used to phosphorylate OmpR1. Increasing concentrations of phosphorylated OmpR1 were incubated with 50 nM DNA containing the *ompR1* promoter. DNA–protein complexes were analyzed by native gel electrophoresis and visualized by GelRed staining. Lane 1, free DNA; lane 2, 0.1 µM; lane 3, 1 µM; lane 4, 2 µM; lane 5, 3 µM; lane 6, 5 µM phosphorylated OmpR1. **(D)** Binding of phosphorylated and non-phosphorylated OmpR1 to the *bvrR* and *virB1* promoter regions. Electrophoretic mobility shift assays were performed using DNA fragments containing the *bvrR* (P_bvrR) or *virB1* (P_virB1) promoter regions. OmpR1 was phosphorylated as described above and incubated with promoter DNA prior to native gel electrophoresis. For each promoter, reactions contained no OmpR1 (–), non-phosphorylated OmpR1 (+), or phosphorylated OmpR1 (∼P). Lanes 1–3 correspond to P_bvrR and lanes 4–6 correspond to P_virB1. Lanes 1 and 4 contained no OmpR1, lanes 2 and 5 contained 0.5 µM OmpR1, and lanes 3 and 6 contained 2.0 µM OmpR1. Phosphorylated OmpR1 produced a prominent mobility shift with the *bvrR* promoter, whereas non-phosphorylated OmpR1 exhibited only weak binding. No detectable binding to the *virB1* promoter was observed under these conditions. **(E)** Autoregulation of *bvrR* and binding of phosphorylated BvrR to the *ompR1*, *bvrR*, *vjbR*, and *virB1* promoter regions. Electrophoretic mobility shift assays were performed using DNA fragments containing the indicated promoter regions. BvrR proteins carrying either N-terminal or C-terminal His tags were phosphorylated by BvrS (0.7 µM) prior to incubation with promoter DNA. DNA–protein complexes were resolved by native gel electrophoresis and visualized by GelRed staining. Arrows indicate shifted DNA–protein complexes. **(E1)** Binding of phosphorylated N-terminal His-tagged BvrR (N-His-BvrR) to the *bvrR* promoter (P_bvrR). Lane 1, no phosphorylated BvrR; lane 2, 0.5 µM; lane 3, 2.0 µM; lane 4, 4.0 µM phosphorylated BvrR. **(E2)** Binding of phosphorylated N-His-BvrR to the *virB1* promoter (P_virB1). Lane 1, no phosphorylated BvrR; lane 2, 0.5 µM; lane 3, 2.0 µM phosphorylated BvrR. **(E3)** Binding of phosphorylated C-terminal His-tagged BvrR (C-His-BvrR) to the *ompR1* (P_ompR1, left three lanes) and *virB1* (P_virB1, right three lanes) promoters. For each promoter, lane 1, no phosphorylated BvrR; lane 2, 0.5 µM; lane 3, 2.0 µM phosphorylated BvrR. **(E4)** Binding of phosphorylated C-His-BvrR to the *vjbR* promoter (P_vjbR). Lane 1, no phosphorylated BvrR; lane 2, 2.0 µM; lane 3, 4.0 µM phosphorylated BvrR. **(F)** Binding of VjbR to the *virB1* and *vjbR* promoter regions. Electrophoretic mobility shift assays were performed using DNA fragments containing the *virB1* (P_virB1) or *vjbR* (P_vjbR) promoter regions. Increasing concentrations of purified VjbR were incubated with 50 nM promoter DNA and analyzed by native gel electrophoresis. Lanes 1–4 correspond to P_virB1 and lanes 5–8 correspond to P_vjbR. For each promoter, lane 1 (lanes 1 and 5), free DNA; lane 2 (lanes 2 and 6), 0.5 µM VjbR; lane 3 (lanes 3 and 7), 2.0 µM VjbR; lane 4 (lanes 4 and 8), 4.0 µM VjbR. Arrows indicate shifted VjbR–DNA complexes. **(G)** In vivo regulation of the *virB1* and *bvrR* promoters by VjbR. Plasmids encoding arabinose-inducible VjbR and NanoLuc (NLuc) reporter fusions to the *virB1* or *bvrR* promoters were co-transformed into *Escherichia coli*. VjbR expression was induced with increasing concentrations of arabinose (0–0.2% [w/v]), and promoter activity was quantified by measuring NLuc-derived relative luminescence units (RLU). Data represent the mean ± SD of n = 3 biological replicates.

### Identification of OmpR1 as a regulator of *Brucella* virulence

The requirement for *ompR1* was unexpected, as this regulator has not previously been implicated in *Brucella* virulence. To further characterize this finding, we examined the repertoire of OmpR-family response regulators encoded by the *B. neotomae* genome and identified nine homologs, including *ompR1* (**S8 Data**). Analysis of the Tn-seq data revealed that *ompR1* was the only homolog that met criteria for conditional requirement for intracellular growth, based on both log₂ fold change and q value. Notably, this group includes *ompR7* (*ctrA*) and *ompR9* (*bvrR*), which have been extensively characterized as key regulators of *Brucella* virulence. Insertions in *bvrR* were depleted in the Tn-seq input pool and are discussed further below [33-36].

### Cross-regulation between the OmpR1/EnvZ1 and BvrR/BvrS two-component systems

To investigate potential cross-regulation between the OmpR1/EnvZ1 and BvrR/BvrS systems in *B. neotomae*, we generated constructs encoding each protein with a 6xHis tag fused to either the N or C terminus, allowing purification and biochemical analysis (**Fig. S9**). The BvrR/BvrS system was included based on its established role in regulating the VirB type IV secretion system and *Brucella* virulence.

EnvZ1 was first autophosphorylated in vitro, after which 3 µM OmpR1 was added and phosphorelay was monitored for up to 2 hours. OmpR1 was rapidly phosphorylated by EnvZ1 within 1 minute, and phosphorylation was maintained throughout the assay period (**Fig. 5B**). We next examined phosphorelay through the BvrR/BvrS system and assessed potential cross-phosphorylation between the two systems. BvrR was efficiently phosphorylated by BvrS within 1 minute, with sustained phosphorylation for up to 2 hours. When OmpR1 was incubated with phosphorylated BvrS, rapid phosphorylation was again observed. In contrast, phosphorylation of BvrR by EnvZ1 was not detected at early time points and became apparent only after approximately 1 hour of incubation. These results indicate that OmpR1 is efficiently phosphorylated by both EnvZ1 and BvrS, whereas BvrR is efficiently phosphorylated only by its cognate kinase BvrS and shows weak, delayed phosphorylation by EnvZ1.

### OmpR1 regulates *bvrR* but not *vjbR* or *virB* directly

Given the evidence for cross-regulation, we next examined whether OmpR1 and BvrR bind to promoter regions of virulence-associated genes. Electrophoretic mobility shift assays (EMSA) demonstrated that phosphorylated OmpR1 binds to its own promoter, indicating autoregulation (**Fig. 5C**). We then tested binding of phosphorylated and non-phosphorylated OmpR1 to the *bvrR* and *virB1* promoters (**Fig. 5D**). Both forms of OmpR1 bound the *bvrR* promoter, with stronger binding observed for the phosphorylated protein. In contrast, neither form bound the *virB1* promoter.

We further tested whether OmpR1 regulates *vjbR*, which is known to control *virB* expression. EMSA assays revealed no binding of either non-phosphorylated or phosphorylated OmpR1 to the *vjbR* promoter (**Fig. S10**). Taken together, these results indicate that OmpR1 regulates *bvrR* expression but does not directly regulate *virB1* or *vjbR*.

### BvrR integrates upstream regulation and directly controls *vjbR* and *virB*

To determine whether BvrR functions downstream of OmpR1, we examined binding of phosphorylated BvrR to promoter regions of virulence-associated genes (**Fig. 5E**). Phosphorylated BvrR bound its own promoter, indicating autoregulation. Both N-terminally and C-terminally His-tagged BvrR variants bound the *bvrR* promoter, indicating that N-terminal His tagging does not abolish DNA binding or autoregulatory activity. However, the N-terminally His-tagged BvrR failed to bind the *virB1* promoter, whereas a C-terminally His-tagged BvrR variant bound the *ompR1*, *vjbR*, and *virB1* promoters. Together, these findings indicate that OmpR1 and BvrR participate in a regulatory relationship, consistent with the phosphorelay results demonstrating cross-regulation between the two systems. The ability of BvrR to bind the *vjbR* and *virB1* promoters further supports a role for BvrR as a direct regulator of these genes.

Given that BvrR regulates *vjbR*, we next examined whether VjbR binds to the *virB1* and *vjbR* promoters (**Fig. 5F**). Purified VjbR bound both promoter regions in vitro. To validate these interactions in vivo, we fused the *virB1* and *bvrR* promoters to an NLuc reporter and modulated VjbR expression using arabinose (**Fig. 5G**). VjbR significantly increased *virB1* expression at 0.05% arabinose, whereas no regulatory activity was observed at the *bvrR* promoter. Taken together, these data indicate that BvrR regulates both *vjbR* and *virB1*, while VjbR specifically regulates *virB1*, consistent with a hierarchical regulatory cascade.

### OmpR1 as an upstream regulator of *Brucella neotomae* virulence

Attempts to generate an in-frame deletion of *bvrR* by allelic exchange were unsuccessful, consistent with prior reports in *Brucella ovis* [37], suggesting that *bvrR* is essential for viability. This interpretation is further supported by Tn-seq data (**Table S1A**), in which *bvrR* (WP_004687763.1) exhibited minimal insertion counts across replicates. To enable functional analysis, a conditional *bvrR* mutant was constructed by expressing *bvrR* under the control of an arabinose-inducible promoter. In the absence of arabinose, the mutant showed no detectable growth in M9 medium with glycerol as the carbon source over 48 h, whereas growth was restored upon addition of 0.02% arabinose (**Fig. S11A**). Similarly, intracellular growth at 48 h post-infection in J774A.1 macrophages was attenuated under limiting arabinose conditions (**Fig. S11B**). These findings support an essential role for *bvrR* in both in vitro growth and intracellular infection.

Based on the biochemical and transcriptional data, we next examined whether OmpR1 functions upstream in the virulence hierarchy. We assessed whether intracellular growth defects of regulatory mutants could be rescued by complementation with downstream regulators. Complementation analysis demonstrated that the intracellular growth defect of the *ompR1* mutant was restored by either OmpR1 or BvrR, but not by VjbR (**Fig. 6A**). In contrast, growth of the conditional *bvrR* mutant was restored only by BvrR, and the *vjbR* mutant was rescued exclusively by VjbR. These results indicate that loss of *ompR1* primarily affects intracellular growth through loss of *bvrR* expression, establishing OmpR1 as a primary upstream regulator.

**Fig. 6.**
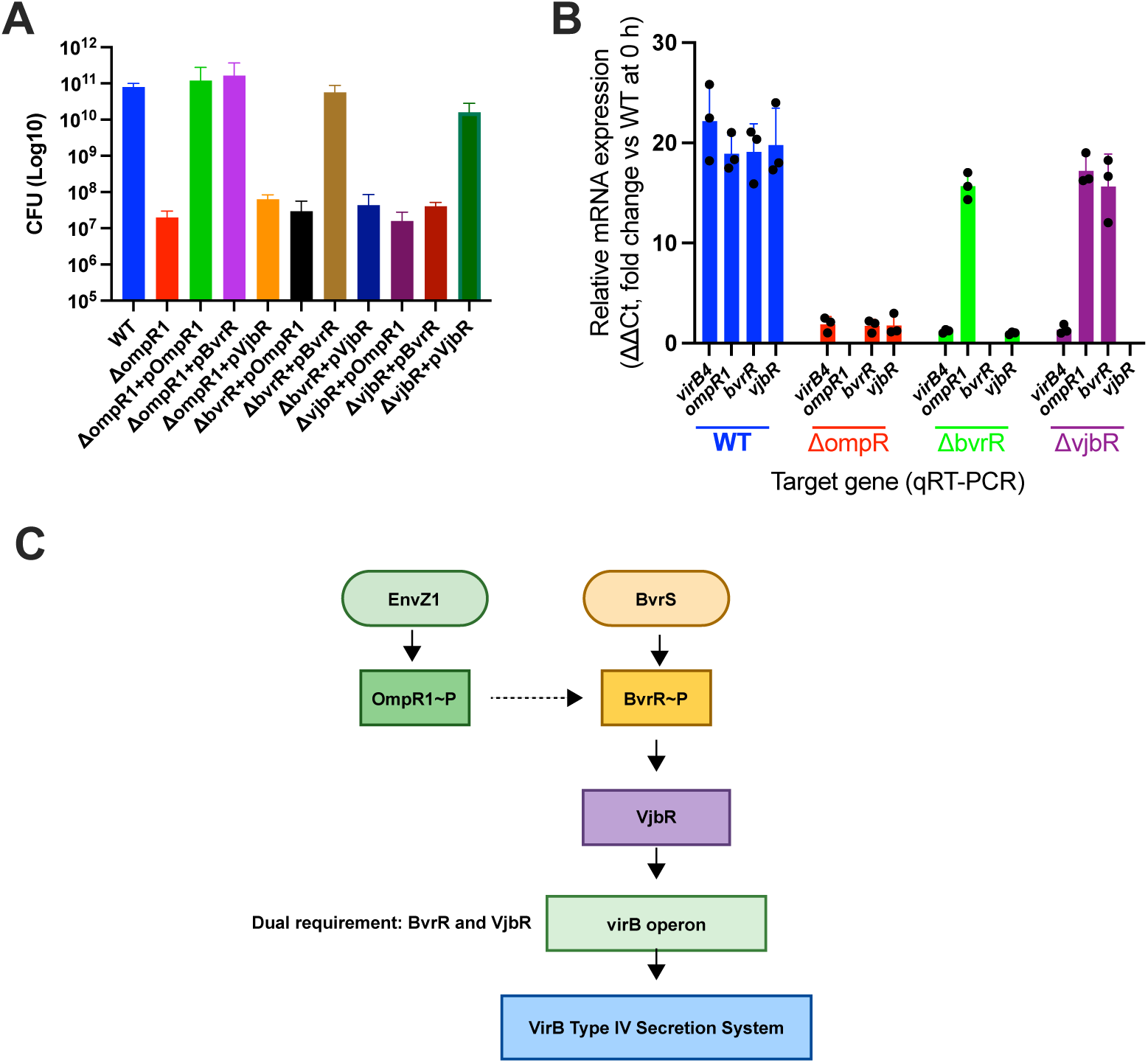
OmpR1 functions as a master regulator of the *Brucella neotomae* virulence circuit. **(A)** Intracellular growth of wild-type (WT) *B. neotomae* and isogenic deletion mutants lacking *ompR1*, *bvrR*, or *vjbR*, along with their corresponding complemented strains, was assessed in J774A.1 macrophages. Cells were infected at an MOI of 10, and intracellular bacterial burdens were quantified by CFU enumeration at 48 h post-infection. Data represent the mean ± SD of n = 3 biological replicates. **(B)** Early infection-stage expression of *virB4*, *ompR1*, *bvrR*, and *vjbR* was measured by qRT-PCR at 4 h post-infection in WT, Δ*ompR1*, Δ*bvrR*, and Δ*vjbR* backgrounds, as indicated by color-coded groupings. Transcript levels were normalized to WT expression at 0 h and are presented as fold change (ΔΔCt). Bars represent the mean ± SD of n = 3 biological replicates, with individual data points overlaid. **(C)** Model of the OmpR1–BvrR–VjbR regulatory hierarchy controlling VirB system activation in *B. neotomae*. The EnvZ1/OmpR1 two-component system functions upstream in the virulence regulatory network and promotes activation of the BvrR/BvrS system through induction of *bvrR* expression. Activated BvrR stimulates transcription of *vjbR*, and VjbR directly activates the *virB* operon. Sequential activation of OmpR1, BvrR, and VjbR is required for full induction of the VirB type IV secretion system under the conditions tested, positioning OmpR1 as a primary upstream activator of the virulence regulatory cascade.

This hierarchy was further supported by transcriptional profiling during early infection. In wild-type *B. neotomae*, high expression of *virB4*, *ompR1*, *bvrR*, and *vjbR* was observed at 4 hours post-infection (**Fig. 6B**). In the *ompR1* mutant, expression of *bvrR*, *vjbR*, and *virB4* was abolished. In the conditional *bvrR* mutant, *ompR1* expression remained intact, whereas *vjbR* and *virB4* were not induced. In the *vjbR* mutant, only *virB4* expression was lost, whereas *ompR1* and *bvrR* expression remained high.

Collectively, these findings position OmpR1 as a primary upstream regulator of the *Brucella neotomae* virulence cascade. These data support a regulatory model in which the OmpR1/EnvZ1 system acts upstream of the BvrR/BvrS system (**Fig. 6C**), with BvrR regulating *vjbR* and *virB1*, and VjbR subsequently contributing to activation of the VirB secretion system.

### Phylogenetic analysis of OmpR1

To determine whether OmpR1, identified here as a regulator of virulence in *B. neotomae*, is conserved across the genus, we identified 119 homologs with high sequence similarity to the *B. neotomae* OmpR1 protein. Phylogenetic analysis (**Fig. S12**) revealed that well-characterized transcriptional regulators, including BvrR (blue clade), PhoB (green clade), and CtrA (magenta clade), cluster tightly with their respective orthologs across the genus. In contrast, OmpR1 forms a distinct lineage relative to these regulatory systems.

Notably, several OmpR1 homologs from other *Brucella* species exhibit near-complete sequence identity to the *Brucella neotomae* protein. Despite this high degree of conservation, these homologs have not been implicated as regulators of virulence in *Brucella*. This apparent absence may reflect limited functional investigation rather than true regulatory divergence, as OmpR-like regulators have not been systematically examined in most species.

## Discussion

In this study, we used genome-wide transposon sequencing to define the genetic determinants required for intracellular survival of *Brucella neotomae* within macrophages. Our findings reveal a requirement for amino acid biosynthesis and a hierarchical regulatory cascade that governs virulence gene expression. These results define key metabolic and regulatory features required for intracellular growth that are not apparent from studies of axenic growth or individual virulence factors.

A prominent feature of the Tn-seq dataset was the enrichment of genes involved in amino acid biosynthesis. Methionine and histidine biosynthesis were required for sustained intracellular growth, while tryptophan contributed partially. Importantly, these requirements were specific to the intracellular environment, as mutants defective in these pathways grew normally in defined minimal medium. These observations indicate that the intracellular growth defects cannot be explained solely by auxotrophy, but instead reflect constraints specific to the intracellular environment.

Previous work has shown that *Brucella* experience nutrient limitation within macrophages, and that amino acid biosynthesis pathways can contribute to intracellular survival. However, reported requirements vary by species and experimental system. For example, methionine biosynthesis mutants of *B. melitensis* and *B. suis* do not exhibit pronounced intracellular growth defects in macrophage models, whereas histidine and other amino acid auxotrophs have been shown to impair intracellular growth in some contexts [26, 27, 38-40]. Consistent with these observations, Tn-seq analyses have reported that genes involved in histidine and leucine biosynthesis exert minimal effects during early stages of infection but become important by 24 hours post-infection [41]. In contrast, our data indicate that *B. neotomae* exhibits a pronounced dependence on amino acid biosynthesis for intracellular growth, including methionine and histidine, with tryptophan contributing partially. Together, these findings indicate that requirements for amino acid biosynthesis during intracellular growth vary across *Brucella* species and between experimental systems.

Amino acid biosynthesis contributes to intracellular growth rather than early virulence gene induction. Supplementation experiments showed that methionine and histidine support intracellular replication in a concentration-dependent manner. Consistent with this, disruption of amino acid biosynthesis did not impair *virB* expression. Instead, expression of biosynthetic genes increased following *virB* induction and was absent in a *virB* mutant background. These observations indicate that amino acid biosynthesis is linked to intracellular growth rather than to early virulence activation, reinforcing the concept that *Brucella* virulence is not a single-stage program but involves distinct phases of gene expression and physiological adaptation.

Beyond metabolism, our analysis identified several regulatory genes required for intracellular survival. Among these, the identification of *ompR1* was unexpected. Although OmpR-family regulators are widespread in bacteria, *ompR1* has not previously been implicated in *Brucella* virulence. Through genetic, biochemical, and transcriptional analyses, we demonstrate that OmpR1 functions at the top of a hierarchical virulence cascade. Specifically, OmpR1 activates the BvrR/BvrS system, which in turn directly regulates *vjbR* and the *virB* operon. Loss of *ompR1* abolishes intracellular growth and cannot be compensated by downstream regulators, supporting a model in which OmpR1 functions upstream in the virulence regulatory hierarchy.

These findings further define a regulatory architecture in which the OmpR1/EnvZ1 system functions upstream of the BvrR/BvrS system to control virulence gene expression. Although cross-phosphorylation between these systems was observed, BvrR-dependent regulation was not observed in the absence of OmpR1, indicating that upstream activation is required for proper signaling through this pathway. In this model, activated BvrR induces *vjbR* and *virB1*, while VjbR further contributes to activation of the VirB secretion system. The apparent lack of feedback from VjbR to upstream regulators suggests a unidirectional regulatory cascade. These observations support a model in which coordinated activation of BvrR and VjbR is necessary for efficient induction of virulence gene expression during intracellular infection.

The identification of OmpR1 as an upstream regulator also provides new context for understanding the role of BvrR/BvrS, which has long been recognized as a central virulence regulator in *Brucella*. Our data suggest that BvrR/BvrS does not function as an independent master regulator, but rather as an intermediate whose activity depends on OmpR1. This distinction may help explain differences observed across species and experimental systems, where BvrR/BvrS-dependent phenotypes vary in magnitude and timing.

Phylogenetic analysis supports the identification of OmpR1 as a distinct group of response regulators. OmpR1 clusters separately from well-characterized *Brucella* response regulators such as BvrR, PhoB, and CtrA. Notably, OmpR1 homologs among species exhibit high sequence conservation yet have not been implicated in virulence. This discrepancy between sequence conservation and functional characterization likely reflects limited functional investigation, rather than true divergence in regulatory role. Together, these findings indicate that OmpR1 is a conserved regulator whose role in intracellular virulence has not been fully explored across the genus.

In this context, our findings suggest that OmpR1 represents a previously uncharacterized regulatory element within the genus, whose role was revealed through unbiased genetic analysis in *B. neotomae*. Evolutionary theory predicts that transcriptional regulators and their target networks can undergo substantial remodeling without extensive changes in protein sequence, particularly through alterations in regulatory connectivity and transcriptional output [42-44]. Whether OmpR1 homologs in other *Brucella* species retain latent regulatory potential, participate in distinct physiological processes, or similarly coordinate virulence programs remains unknown. Addressing these possibilities will require systematic functional interrogation across the genus.

Beyond transcriptional control, our dataset also highlights additional functional classes that support intracellular growth. In particular, we validated the predicted aquaporin gene *aqpZ* as a transport-associated determinant. Deletion of *aqpZ* did not impair growth in rich or minimal media under standard conditions, yet produced a marked defect in macrophage replication. The Δ*aqpZ* mutant also exhibited increased sensitivity to elevated NaCl, consistent with prior studies in *Brucella abortus* demonstrating induction of aquaporin expression under hyperosmotic conditions and a role in osmotolerance [45]. However, a contribution of aquaporins to intracellular replication has not previously been established. Although the physicochemical properties of the *Brucella*-containing vacuole remain incompletely defined, these findings indicate that osmotic stress resistance contributes to growth during host infection. The mechanistic basis linking these phenotypes warrants further investigation.

Several limitations of this study should be acknowledged. Our experiments were performed in a macrophage cell line model and focused on a single *Brucella* species. Whether the regulatory architecture defined here operates identically in primary macrophages, in vivo infection models, or under different intracellular conditions remains to be determined. In addition, although we establish OmpR1 at the apex of the virulence hierarchy, the upstream signals sensed by the OmpR1/EnvZ1 system are not yet defined. Identifying these cues will be critical for understanding how *Brucella* detects host-derived signals and initiates intracellular adaptation.

In summary, this work identifies genetic determinants required for intracellular growth in *B. neotomae* and establishes OmpR1 as a central regulator of virulence gene expression. By combining genome-wide fitness profiling with targeted functional analyses, we show that metabolic capacity, regulatory control, and osmotic homeostasis support intracellular replication. These findings provide a framework for understanding how *Brucella* adapts to the intracellular environment and suggest directions for examining related pathways across the genus.

## Materials & Methods

### Bacterial strains, cell lines, and culture conditions

Bacterial strains and eukaryotic cell lines used in this study are listed in Table S13. *Escherichia coli* NEB-5α (New England Biolabs, Ipswich, MA), Ec100D pir-116 (Bioresearch Technologies, Petaluma, CA), and β2155 [46] were grown at 37 °C with shaking in Luria broth (LB) (BD, Franklin Lakes, NJ). *Brucella neotomae* (Bn) 5K33 WT, Bn-Lux, Bn-tdTomato, and Bn Δ*virB4* [21] were used for strain construction and experimental work. *Brucella* strains were grown at 37 °C in trypticase soy broth (TSB) (BD) or M9 minimal medium in a humidified incubator with 5% CO₂.

Plasmid isolation, transformation, and PCR amplification were performed as previously described [21]. *E. coli* and *B. neotomae* strains were grown in the presence of 50 µg/mL nourseothricin (Ntc), 50 µg/mL kanamycin (Km), 10 µg/mL chloramphenicol (Cam), or 100 µg/mL ampicillin (Amp), as appropriate. Diaminopimelic acid (DAP; 50 µg/mL) was added for growth of β2155, and TSB was supplemented with 10% sucrose for sacB counterselection during construction of *Brucella* deletion mutants.

Murine J774A.1 (ATCC TIB-67) macrophages were maintained in RPMI 1640 (Corning, Corning, NY) supplemented with 9% heat-inactivated, iron-supplemented calf serum (GemCell, Gemini Bio-Products, West Sacramento, CA) at 37 °C in a humidified atmosphere containing 5% CO₂.

### Construction of *B. neotomae* transposon pool

A modified Himar1 transposon delivery system was developed using the pMar2xT7 vector [47] as a backbone. To enable antibiotic selection in the target organism, a nourseothricin resistance gene (NAT) was substituted for the gentamicin resistance cassette between the inverted repeats (IRs) of the transposon. In addition, the native IR sequence (GCCAAC) was mutated to TCCGAC to introduce a recognition site for the MmeI restriction enzyme, facilitating downstream insertion site analysis. The resulting plasmid, designated pMar-Nat, was transformed into the diaminopimelic acid (DAP) auxotroph *E. coli* β2155 by electroporation.

The transposon library was generated in the Bn WT strain (type strain 5K33) by tri-parental conjugation. Donor (*E. coli* β2155 carrying pMar-Nat), recipient (Bn WT), and helper (*E. coli* carrying the RK600 plasmid) strains were mixed at a ratio of 1:2:1. The mixture was resuspended and spotted onto tryptic soy agar (TSA) plates supplemented with 50 µg/mL DAP and incubated at 37 °C for 24 hours. Following incubation, the bacterial lawn was harvested and resuspended in sterile PBS. The suspension was plated onto TSA containing 50 µg/mL nourseothricin to select for transconjugants. Using this protocol, a high-density transposon library comprising approximately 150,000 mutants was obtained. The B. neotomae Tn-seq library was supplemented with sterile glycerol to a final concentration of 50% (v/v), aliquoted, and stored at −80 °C.

### Tn-seq Analysis

To identify *B. neotomae* genes required for intracellular survival, 1 × 10⁷ J774A.1 macrophages were seeded in 10-cm petri dishes and allowed to adhere for 24 hours. Cells were then infected with the *B. neotomae* mutant library at a multiplicity of infection (MOI) of 10 to maximize representation of the mutant library during infection of the host cell population. Under these conditions, prior analysis indicates that most infected macrophages harbor one or few bacteria with limited colocalization [21]. At 4 hours post-infection, gentamicin (100 µg/mL) was added to the plates and incubated for 1 hour to kill extracellular bacteria. Cells were then washed three times with sterile DPBS to remove non-adherent bacteria and residual antibiotic and maintained in fresh RPMI medium supplemented with gentamicin (10 µg/mL) to prevent outgrowth of extracellular bacteria.

Genomic DNA (gDNA) was extracted from the initial library (input pool) used for infection. At 48 hours post-infection, infected macrophages were incubated in 0.2% saponin on ice for 30 minutes to permeabilize host cell membranes while preserving bacterial integrity. Cells were then scraped to detach adherent material and combined with the supernatant. The resulting suspension, containing bacteria and host cell debris, was pelleted by centrifugation at 3,500 × g for 10 minutes at 4 °C. Genomic DNA was extracted from the pellet using the Promega Wizard genomic DNA purification kit according to the manufacturer’s instructions.

Genomic DNA samples from both pools were digested with MmeI. Resulting fragments were ligated to adapters containing 6-bp barcodes (Table S1): input pool (three replicates; CGATGT, ATCACG, and TTAGGC) and output pool (three replicates; TGACCA, ACAGTG, and GCCAAT). Transposon–genome junctions were selectively amplified by PCR using Tn-seq-F/R primers (**Table S1**). Six PCR-amplified samples (three input and three output replicates) were submitted to the TUCF Genomics Core for Illumina sequencing.

The raw sequencing data (FASTQ files) were processed using the Galaxy platform [48]. Reads were trimmed to remove the 27-bp inverted repeat (IR) sequence (5′-AGACCGGGGACTTATCATCCGACCTGT), leaving 16-bp chromosomal sequences flanking insertion sites for downstream analysis. Trimmed reads were aligned to the *Brucella neotomae* reference genome (RefSeq assembly GCF_000742255.1; nucleotide accession NZ_KN046827.1) to identify transposon–chromosome junctions, and alignments were exported as SAM (Sequence Alignment Map) files. These files were analyzed using Tn-seq Explorer [49] to quantify transposon insertions per gene based on the mapping positions of the 16-bp reads for both input and output pools.

### Construction of *B. neotomae* mutants and complementary strains

To delete genes identified as putatively essential by Tn-seq analysis, in-frame internal deletions were generated using an overlapping PCR strategy. For each target gene, two fragments were amplified: one containing the upstream flanking region and the 5′ terminal portion of the coding sequence, and the other containing the 3′ terminal portion of the coding sequence and the downstream flanking region. Fragments were purified using a PCR clean-up kit (Qiagen) and joined by overlap PCR to generate constructs lacking the internal coding region while retaining short N- and C-terminal coding sequences (approximately 5–10 amino acids). This approach minimizes disruption of translational coupling and preserves regulatory elements near the 3′ end of the gene.

The resulting PCR product was digested with BamHI and SacI and ligated into the suicide vector pSR47S [50], which requires the λ Pir protein provided in trans for replication. Recombinant plasmids were transformed into *E. coli* Ec100D pir⁺ for propagation and sequence verification. Sequence-verified constructs were introduced into Bn-Lux or Bn-tdTomato strains by electroporation (1.8 kV, 400 Ω, 25 µF). Cells were recovered in TSB for 1 h at 37 °C and plated on TSA containing 50 µg/mL kanamycin to select for single-crossover integrants. Integrants were subsequently plated on TSA supplemented with 10% sucrose for counterselection. Candidate colonies were screened by colony PCR using gene-specific 1F and 2R primers to identify clones with deletion of the internal coding region.

To complement gene deletions in *B. neotomae* strains, complete coding sequences of the target genes were amplified using gene-specific complementation primers (**Table S13**). Amplicons were digested and ligated into pBMTL-2 [51] at the XbaI and EcoRV restriction sites. Recombinant pBMTL-2 plasmids were introduced into the corresponding *B. neotomae* mutant strains by electroporation. Cells were recovered in 1 mL TSB for 1 h at 37 °C and plated on TSA supplemented with 50 µg/mL kanamycin to select for complemented strains.

### *B. neotomae* infection and intracellular growth

To assess intracellular survival of *B. neotomae* strains, J774A.1 murine macrophages were seeded at a density of 5 × 10⁴ cells per well in 96-well plates and incubated at 37 °C with 5% CO₂. After 24 hours, macrophages were infected with Bn-tdTomato or Bn-Lux strains (wild-type, deletion mutants, or complemented strains) at a multiplicity of infection (MOI) of 10. To synchronize infection, plates were centrifuged at 930 × g for 10 minutes. At 4 hours post-infection, extracellular bacteria were killed by treatment with gentamicin (100 µg/mL) for 1 hour. Wells were then washed twice with PBS, and fresh RPMI medium was added.

Intracellular bacterial growth was assessed using two complementary approaches. For strains expressing luciferase (Bn-Lux), intracellular growth was measured by luminescence at specified time points using an Infinite M1000 PRO plate reader. For strains labeled with tdTomato, intracellular bacterial burden was quantified by CFU enumeration at 48 h post-infection. Macrophages were lysed, and serial dilutions were plated on TSB agar supplemented with nourseothricin to determine colony-forming units (CFU). The tdTomato-nat cassette was used for selection and was not used as an alternative quantitative readout.

### Protein Expression and Purification

For protein expression of OmpR1/EnvZ1, BvrR/BvrS, and VjbR, recombinant plasmids were designed to incorporate a 6x His tag at either the N-terminus or C-terminus of each protein. Briefly, the coding sequences of the target genes were amplified by PCR using gene-specific expression primers (**Table S13**). For N-terminal His-tagging, amplicons were digested and ligated into the pBAD vector at the BglII and EcoRI restriction sites. For C-terminal His-tagging of BvrR, the gene was inserted into the pBAD backbone using NcoI and EcoRI sites. The resulting constructs were transformed into *E. coli* NEB-5α competent cells for propagation and sequence verification.

To express the recombinant proteins, an overnight culture of NEB-5α was inoculated at a 1:100 ratio into 10 mL of fresh LB medium supplemented with 100 μg/mL ampicillin. The culture was incubated at 37°C for 5 hours, after which protein expression was induced by the addition of L-arabinose to a final concentration of 0.2%. Following overnight incubation, the cells were harvested by centrifugation at 4,000 × g for 10 minutes. The resulting cell pellet was resuspended in 900 μL of lysis buffer (50 mM Tris-Cl, 50 mM NaCl, pH 7.5) supplemented with 100 μL of lysozyme solution (10 mg/mL), 1 mM PMSF, and 3 U/mL Benzonase nuclease (Sigma-Aldrich). For EnvZ1 and BvrS, the buffer was additionally supplemented with 3% n-dodecyl-β-D-maltoside (DDM). The suspension was chilled on ice and lysed by ultrasonication using a 1/8-inch microtip probe (50% amplitude; 5 cycles of 40 s on and 59 s off) with a Fisherbrand Model 120 Sonic Dismembrator. The lysate was then centrifuged at 12,000 × g for 15 minutes at 4°C, and the supernatant (soluble fraction) was collected for purification.

Protein purification was performed using Ni-NTA spin columns (Qiagen, Valencia, CA). The columns were first equilibrated with lysis buffer containing 10 mM imidazole. The soluble lysate was loaded onto the column and centrifuged at 900 × g for 5 minutes; this loading step was repeated to ensure maximum binding. The column was washed with buffer containing 20 mM imidazole at 900 × g for 2 minutes to remove non-specific proteins. The target proteins were eluted twice using an elution buffer containing 500 mM imidazole. The lysate (L), wash (W), and final eluate (E) fractions were mixed with sample buffer and boiled for 5 minutes to denature the proteins. The prepared samples were loaded onto an SDS-PAGE gel. Following electrophoresis, the gel was immersed in staining buffer for 2 hours, followed by incubation in destaining buffer until protein bands were clearly visible. For downstream applications, the purified proteins were dialyzed against a buffer of 50 mM Tris-Cl and 50 mM NaCl (pH 7.5) for 24 h to remove imidazole. The final protein concentration was determined using a BCA protein assay kit (ThermoFisher Scientific).

### Construction and testing of promoter probe vectors

To create a promoterless reporter backbone, the NLuc gene [52] was amplified and ligated into the pBMTL3 vector at the XbaI and HindIII restriction sites. The recombinant plasmid was transformed into *E. coli* NEB-5α competent cells. Subsequently, DNA fragments containing the promoter regions of *virB1* or *bvrR* were amplified and inserted into the BamHI and XbaI sites upstream of the NLuc gene in the pBMTL3-NLuc vector, generating the reporter constructs P_virB1-NLuc and P_bvrR-NLuc. Each reporter plasmid was co-transformed with the pBAD-VjbR expression vector into *E. coli* NEB-5α cells. Transformants harboring both plasmids were selected on LB agar supplemented with chloramphenicol (10 μg/mL) and ampicillin (100 μg/mL). To induce VjbR expression, cultures were treated with increasing concentrations of L-arabinose (0–0.2%). NLuc activity was quantified by addition of furimazine followed by measurement of luminescence, allowing assessment of VjbR-dependent regulation of the target promoters.

### Autophosphorylation and phosphorelay assays

Autophosphorylation of EnvZ1 and BvrS was performed as follows. Briefly, reactions were conducted in buffer containing 50 mM Tris-Cl (pH 7.5), 50 mM KCl, 10 mM MgCl₂, and 2 mM DTT. Assays were initiated by adding 50 μM ATP to 0.7 μM EnvZ1 or BvrS in a total volume of 20 μL and incubated at room temperature for 1 h. Phosphorylated proteins were used directly in phosphorelay assays.

To investigate phosphotransfer within the OmpR1/EnvZ1, BvrR/BvrS, OmpR1/BvrS, and BvrR/EnvZ1 systems, reactions were initiated by adding 20 μL of autophosphorylated EnvZ1 or BvrS to mixtures containing the cognate response regulator (3 μM OmpR1 or 4 μM BvrR). The final volume was adjusted to 50 μL in kinase buffer (50 mM Tris-Cl [pH 7.5], 50 mM KCl, 10 mM MgCl₂, and 2 mM DTT). Reactions were incubated at room temperature for 0–2 h. At the indicated time points, reactions were terminated by addition of 5× SDS-PAGE sample buffer. Samples were resolved on Phos-tag SDS-PAGE gels (FUJIFILM Wako Chemicals, Richmond, VA) to separate phosphorylated (OmpR1∼P or BvrR∼P) and unphosphorylated species. Gels were stained with Coomassie Brilliant Blue R-250 for 1 h, destained, and imaged using a Bio-Rad ChemiDoc system.

### Protein-promoter interactions

To evaluate DNA binding by OmpR1 and BvrR under phosphorylating conditions, phosphotransfer was coupled to the EMSA reaction. EnvZ1 or BvrS was first autophosphorylated, after which 0.7 µM of the activated kinase was incubated with OmpR1 or BvrR at the indicated concentrations to allow phosphotransfer. Based on prior phosphorelay assays, the reaction was allowed to proceed for 5 min before addition of 50 nM target promoter DNA (**Table S13**). DNA binding was then assessed in EMSA binding buffer (100 mM HEPES, 100 mM KCl, 1 mM DTT, 25% glycerol). Reactions were incubated for 1 h at room temperature. For VjbR, protein was incubated directly with promoter DNA in the same EMSA buffer without kinase pre-treatment. Reactions were terminated by addition of native 5× loading buffer and resolved on Mini-PROTEAN TGX precast polyacrylamide gels (Bio-Rad) in 1× TBE at 100 V for 1 h. Gels were stained in TE buffer containing 1× GelRed for 30 min, and DNA mobility shifts were visualized using a Bio-Rad GelDoc imaging system.

### RNA extraction, cDNA synthesis and real-time PCR reactions

To analyze transcriptional profiles of *B. neotomae* WT and mutant strains during infection, J774A.1 macrophages were infected at an MOI of 10. At indicated time points postinfection, macrophages were harvested and lysed to recover intracellular bacteria. Briefly, cells were incubated in 0.2% saponin on ice for 30 min to lyse host cells while maintaining bacterial integrity. Bacterial pellets were collected by centrifugation at 3,500 × g for 10 min at 4°C. Total RNA was extracted using the Direct-zol RNA Miniprep Kit (Zymo Research, Irvine, CA) according to the manufacturer’s instructions, including on-column DNase I treatment to remove genomic DNA. RNA concentration and purity were assessed by measuring absorbance at 260 and 280 nm. Subsequently, 1 μg of total RNA was reverse-transcribed into cDNA using ProtoScript II Reverse Transcriptase (NEB, Beverly, MA).

The resulting cDNA was used as a template for quantitative PCR (qPCR) to measure expression of target genes. Reactions were performed using the DyNAmo HS SYBR Green qPCR Kit (Thermo Scientific) on a Bio-Rad CFX384 Real-Time PCR Detection System. Each reaction contained 1 µL of cDNA in a total volume according to the manufacturer’s protocol. Gene expression was normalized to *rpoB* as an internal reference. Relative expression levels were calculated using the ΔΔCt method and used to compare transcript abundance between wild-type and mutant strains.

## Supporting information

S2_Table_TnSeq_statistics

S3_Table_Volcano_stats

S4_Data_Raw_Figure_Data

S8_Data_OmpR1_homologs_fasta

S13_Table_Strains_Primers

## Funding and Acknowledgements

This work was supported by the National Institute of Allergy and Infectious Diseases of the National Institutes of Health under award number R01AI099122 to J.E.K. and by a Novel Therapeutics Development Grant from the Massachusetts Life Sciences Center. The content is solely the responsibility of the authors and does not necessarily represent the official views of the National Institutes of Health. We thank TECAN for use of the Infinite M1000 Pro and D300 instrumentation. TECAN was not involved in the design of the study, preparation of the manuscript, or the decision to submit for publication.

## Supporting Information

Table S1. Adapters and primers used for Tn-seq library construction. Oligonucleotide sequences used for Himar1 transposon library preparation and sequencing.

Table S2. Genome-wide transposon insertion statistics for *Brucella neotomae*. Genome-wide Tn-seq data generated from input and intracellular output pools across three independent biological replicates. Separate tabs provide: (A) per-gene insertion counts and calculated fitness statistics, (B) genes significantly depleted following intracellular growth (FDR ≤ 10%), and (C) genes exhibiting strong depletion (log2 fold change ≥ 2.5).

Table S3. Gene-level Tn-seq fitness statistics used for volcano plot analysis. Complete gene-level statistical output including log2 fold change, p values, and FDR-adjusted q values used to generate the volcano plot shown in Fig. 2.

S4 Data. Raw numerical data underlying Figs. 3-6 and Figs. S5-S7 and S10. Excel file containing raw numerical values underlying quantitative graphs in the main and selected supplemental figures. Separate tabs correspond to individual figure panels as labeled.

Fig. S5. Intracellular growth of histidine biosynthesis mutants.

Fig. S6. Growth of *B. neotomae* strains in minimal medium.

Fig. S7. Deletion of *aqpZ* impairs intracellular replication and osmotic stress tolerance.

S8 Data. OmpR homolog protein sequences (FASTA format). FASTA-formatted amino acid sequences used for phylogenetic analysis shown in Fig. S10.

Fig. S9. Purification of His-tagged recombinant proteins.

Fig. S10. OmpR1 does not bind the *vjbR* promoter.

Fig. S11. Conditional depletion of *bvrR* impairs growth in vitro and during intracellular infection.

Fig. S12. Phylogenetic analysis of OmpR1 and related response regulators.

Table S13. Bacterial strains, plasmids, cell lines, and PCR primers used in this study. Comprehensive list of strains, plasmids, expression constructs, and PCR primers used for mutant construction, complementation, promoter analysis, and gene expression studies.

