## Supplementary material for "Genome-wide identification of metabolic and regulatory determinants of intracellular growth in *Brucella neotomae*": S1_Table

**Table S1. Adapters and primers used for Tn-seq library construction**

| Description | Sequences (5’-3’) | Usage |
| --- | --- | --- |
| BnAD_1A | TTCCCTACACGACGCTCTTCCGATCT**CGATGT**NN | Adapter having CGATGT barcode |
| BnAD_1B | 5Phos/ **ACATCG**AGATCGGAAGAGCGTCGTGTAGGGAAAGAG /3Phos |  |
| BnAD_2A | TTCCCTACACGACGCTCTTCCGATCT**ATCACG**NN | Adapter having ATCACG barcode |
| BnAD_2B | 5Phos/ **CGTGAT**AGATCGGAAGAGCGTCGTGTAGGGAAAGAG /3Phos |  |
| BnAD_3A | TTCCCTACACGACGCTCTTCCGATCT**TTAGGC**NN | Adapter having TTAGGC barcode |
| BnAD_3B | 5Phos/ **GCCTAA**AGATCGGAAGAGCGTCGTGTAGGGAAAGAG /3Phos |  |
| BnAD_4A | TTCCCTACACGACGCTCTTCCGATCT**TGACCA**NN | Adapter having TGACCA barcode |
| BnAD_4B | 5Phos/ **TGGTCA**AGATCGGAAGAGCGTCGTGTAGGGAAAGAG /3Phos |  |
| BnAD_5A | TTCCCTACACGACGCTCTTCCGATCT**ACAGTG**NN | Adapter having ACAGTG barcode |
| BnAD_5B | 5Phos/ **CACTGT**AGATCGGAAGAGCGTCGTGTAGGGAAAGAG /3Phos |  |
| BnAD_6A | TTCCCTACACGACGCTCTTCCGATCT**GCCAAT**NN | Adapter having GCCAAT barcode |
| BnAD_6B | 5Phos/ **ATTGGC**AGATCGGAAGAGCGTCGTGTAGGGAAAGAG /3Phos |  |
| Tn-F | AATGATACGGCGACCACCGAGATCACACTCTTTCCCTACACGACGCTCTTCCGATCT (Adapter) | Amplification of Bn genomic DNA produced by MmeI cleavage |
| Tn-R | *CAAGCAGAAGACGGCATACGA*AGACCGGGGACTTATCATCCGACCTGT (Inverted Repeat) |  |
