## Supplementary material for "Genome-wide identification of metabolic and regulatory determinants of intracellular growth in *Brucella neotomae*": S5_Figure

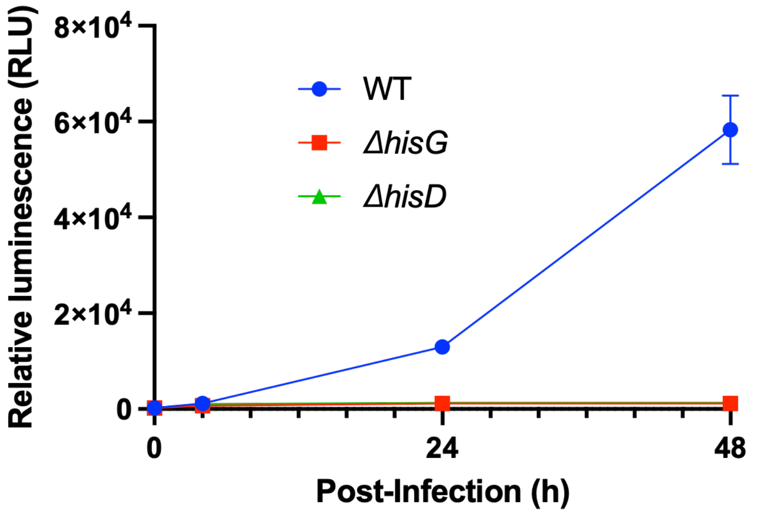


**S5 Figure. Intracellular growth of *B. neotomae* histidine biosynthesis mutants.** Intracellular growth of the wild-type (WT) strain and the Δ*hisG* and Δ*hisD* mutant strains was monitored at 0, 4, 24, and 48 hours post-infection. J774A.1 macrophages were infected at a multiplicity of infection (MOI) of 10. Intracellular bacterial replication was quantified by measuring luminescence emitted from a chromosomally integrated *lux* operon. Data represent the mean of four independent experiments, with standard deviations indicated.
