## Supplementary material for "Genome-wide identification of metabolic and regulatory determinants of intracellular growth in *Brucella neotomae*": S6_Figure

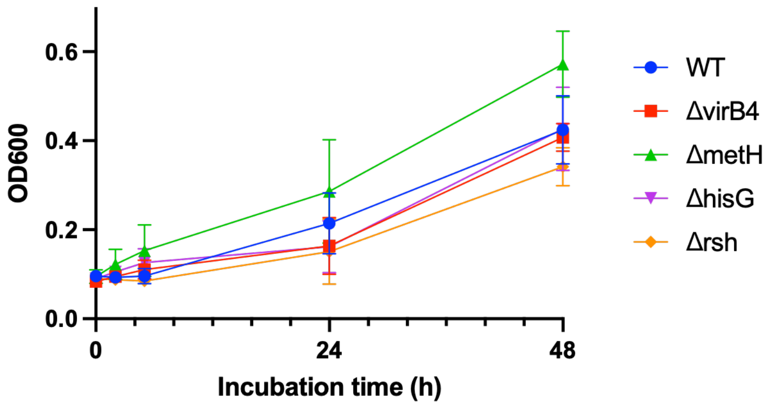


**S6 Figure. Growth of *B. neotomae* strains in minimal medium.** To assess whether intracellular growth defects reflect a general impairment in bacterial proliferation under nutrient-limiting conditions, the wild-type (WT) strain and mutant strains (Δ*virB4*, Δ*metH*, Δ*hisG*, and Δ*rsh*) were cultured in M9 minimal medium. Bacterial growth was monitored by measuring optical density at 600 nm (OD600) at 0, 2, 5, 24, and 48 hours. All mutant strains reached OD600 values comparable to WT by 48 hours, indicating the absence of a generalized growth defect during axenic growth. Data represent the mean ± SD of four independent experiments.
