## Supplementary material for "Genome-wide identification of metabolic and regulatory determinants of intracellular growth in *Brucella neotomae*": S7_Figure

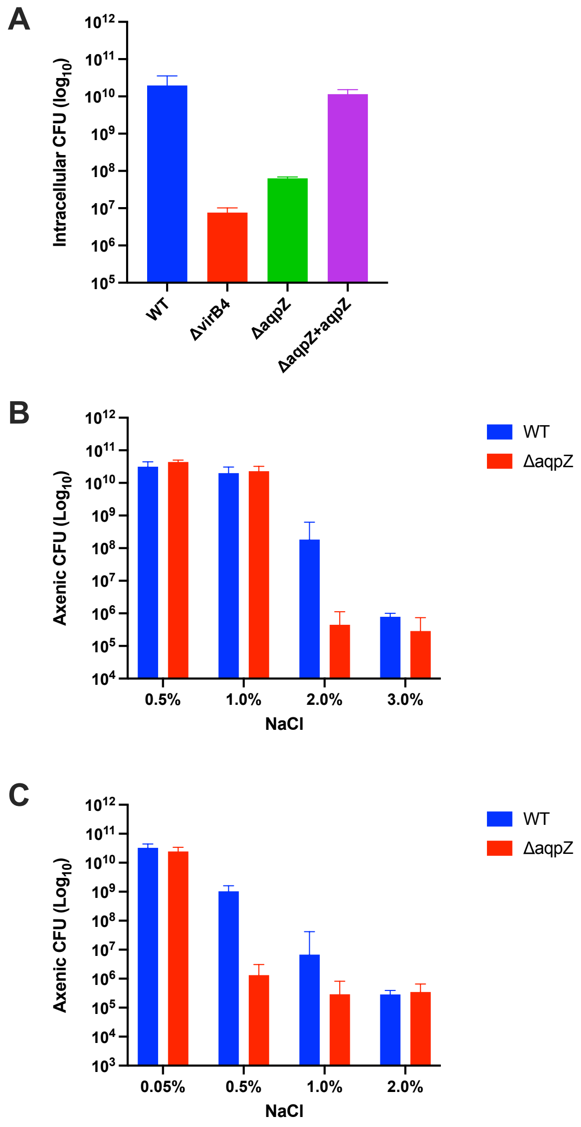


**Supplementary Figure S7. Deletion of *aqpZ* impairs intracellular replication and osmotic stress tolerance. (A)** Intracellular replication of WT, Δ*virB4*, Δ*aqpZ*, and complemented Δ*aqpZ* strains in J774A.1 macrophages. Cells were infected at an MOI of 10, and intracellular CFU were enumerated at 48 h post-infection. The Δ*aqpZ* mutant exhibited a marked intracellular replication defect that was restored by plasmid-based complementation. Δ*virB4* served as a non-replicative control. **(B)** Axenic growth of WT and Δ*aqpZ* strains in TSB supplemented with increasing concentrations of NaCl. Cultures were incubated under standard conditions and CFU were determined at 48 h. No growth defect was observed under standard TSB conditions; however, the Δ*aqpZ* strain displayed increased sensitivity at higher NaCl concentrations. **(C)** Axenic growth of WT and Δ*aqpZ* strains in M9 minimal medium supplemented with increasing concentrations of NaCl. Growth was assessed at 48 h. The Δ*aqpZ* mutant exhibited enhanced salt sensitivity relative to WT, with growth impairment observed at lower NaCl concentrations compared to TSB. Data represent mean ± SD from n = 3 biological replicates.
