## Supplementary material for "Genome-wide identification of metabolic and regulatory determinants of intracellular growth in *Brucella neotomae*": S9_Figure

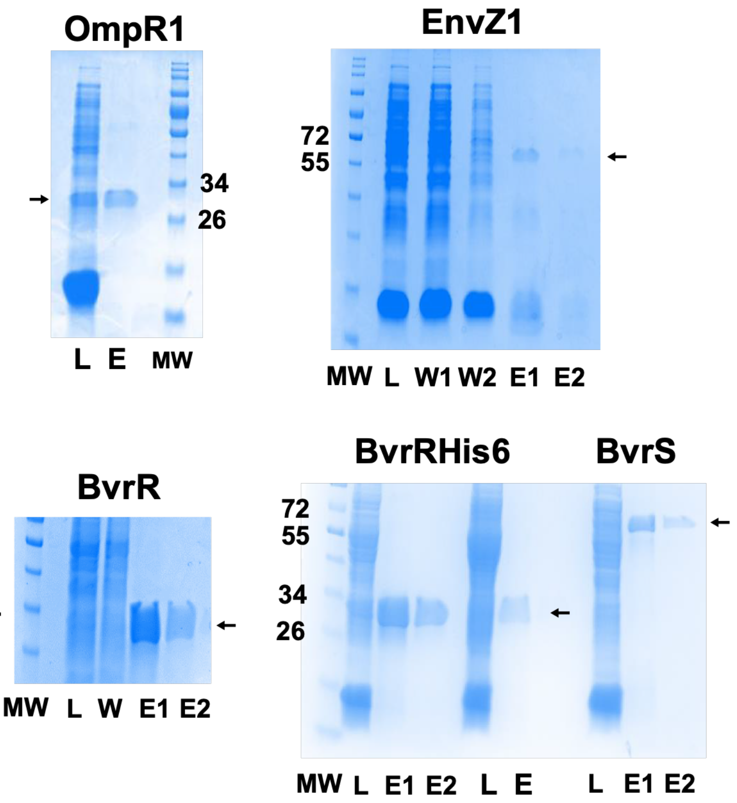


**S8 Figure. Purification of His-tagged Recombinant Proteins for Biochemical Assays.** EnvZ1, BvrR, and BvrS proteins were expressed with a 6×His tag fused to either the N- or C-terminus and purified by Ni-NTA affinity chromatography. Lysate (L), wash (W or W1/W2), and elution (E or E1/E2) fractions were resolved by SDS–PAGE and visualized by Coomassie staining. Molecular weight markers (MW) are indicated. Arrows denote the predominant recombinant protein present in the elution fractions used for downstream EMSA and phosphorelay assays. For BvrR, the N-terminally His-tagged construct is labeled His6-BvrR (bottom left) and the C-terminally His-tagged construct is labeled BvrR-His6 (bottom middle).
