## Supplementary material for "Genome-wide identification of metabolic and regulatory determinants of intracellular growth in *Brucella neotomae*": S10_Figure

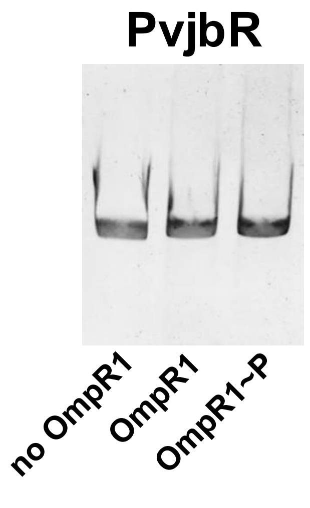


**S9 Figure. OmpR1 does not bind the *vjbR* promoter.** Electrophoretic mobility shift assays (EMSA) were performed to assess binding of OmpR1 to the *vjbR* promoter. A 50 nM DNA fragment corresponding to the *vjbR* promoter was incubated in the absence of protein (lane 1), with non-phosphorylated OmpR1 (lane 2), or with phosphorylated OmpR1 (pOmpR1, lane 3). OmpR1 phosphorylation was achieved using its cognate kinase EnvZ1. Reaction mixtures were resolved on a native polyacrylamide gel and DNA was visualized by GelRed staining. No DNA mobility shift was observed under any condition, indicating that OmpR1 does not directly bind the *vjbR* promoter under these assay conditions.
