## Supplementary material for "Genome-wide identification of metabolic and regulatory determinants of intracellular growth in *Brucella neotomae*": S11_Figure

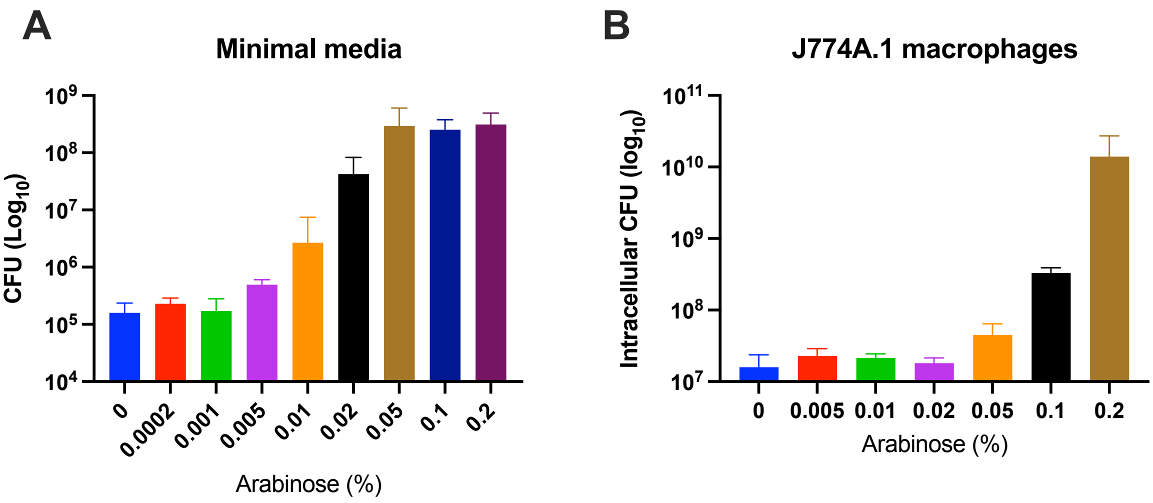


**Figure S11. Conditional depletion of *bvrR* impairs growth in vitro and during intracellular infection.**

(A) Growth of the conditional *bvrR* mutant in M9 minimal medium supplemented with glycerol as the carbon source at increasing concentrations of arabinose. In the absence of arabinose, no detectable growth was observed over 48 h, whereas growth was progressively restored with increasing arabinose concentrations.

(B) Intracellular replication of the conditional *bvrR* mutant in J774A.1 macrophages at 48 h post-infection. Bacterial burden increased in an arabinose-dependent manner, with attenuated growth observed under limiting arabinose conditions.

Data represent the mean ± SD of n = 3 biological replicates.
