## Supplementary material for "Genome-wide identification of metabolic and regulatory determinants of intracellular growth in *Brucella neotomae*": S12_Figure

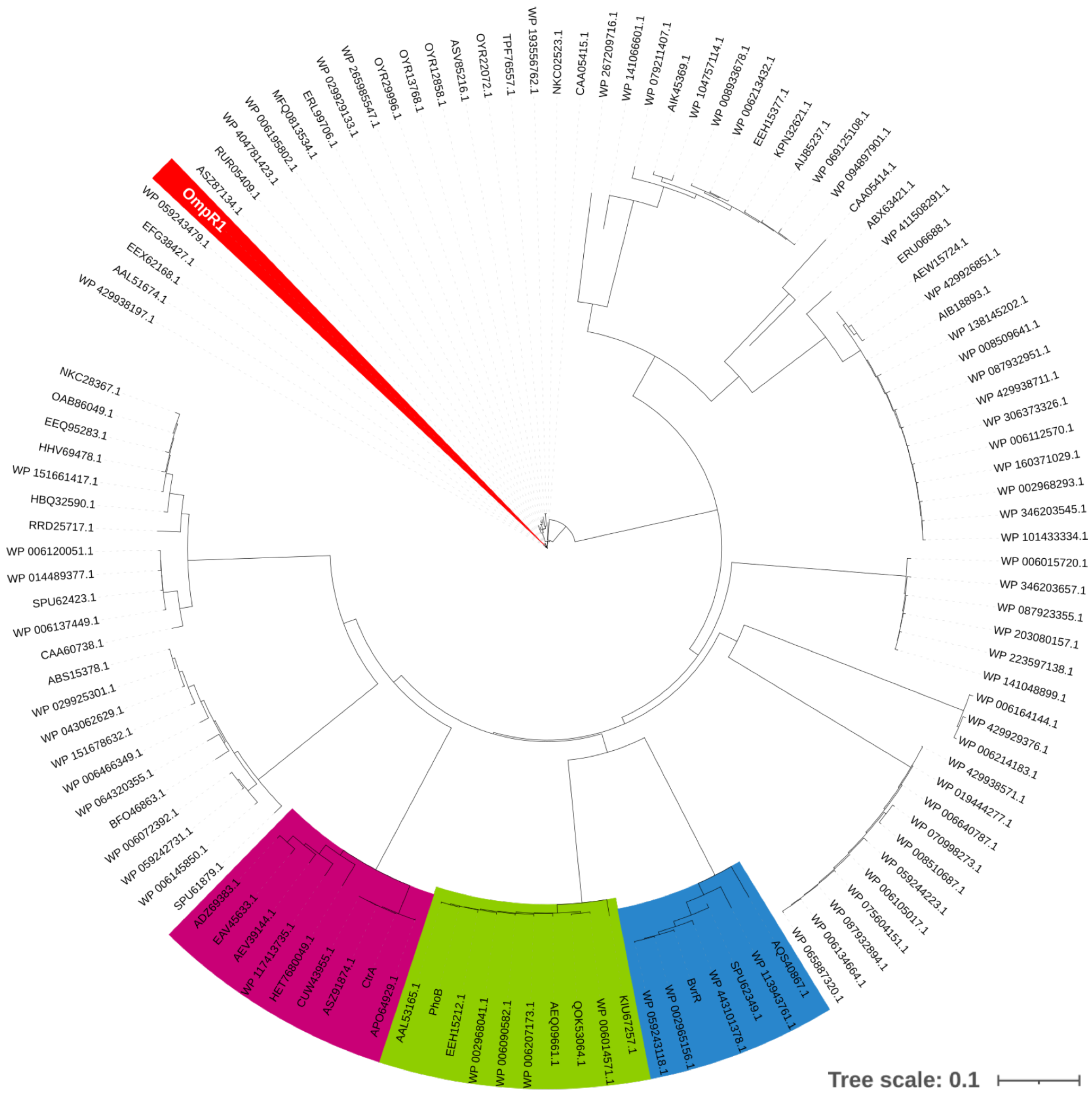


**S10 Figure. Phylogenetic analysis of OmpR1 and related response regulators in Brucella.** A circular phylogenetic tree depicting evolutionary relationships among OmpR1 and representative response regulators within the genus *Brucella*. Protein sequences were aligned using Clustal Omega, and the tree was visualized using iTOL. Canonical response regulators BvrR (blue), PhoB (green), and CtrA (magenta) cluster with their respective orthologs, whereas OmpR1 forms a distinct clade that is clearly separated from these well-characterized regulatory families. Branch lengths are proportional to amino acid substitutions per site (scale bar, 0.1).
