## Supplementary material for "Genome-wide identification of metabolic and regulatory determinants of intracellular growth in *Brucella neotomae*": S13_Table_Strains_Primers

**Table S13. Bacterial strains, plasmids, cell lines, and PCR primers used in this study.**

| **Bacterial strains** | | |
|---|---|---|
| *Strain* | *Relevant characteristics* | *Source or Reference* |
| *B. neotomae* 5K33 | Parent biosafety level 2 rodent pathogen | BEI Resources |
| *B. neotomae*-Lux | Transposon mutant of 5K33 expressing Lux operon-nat genes | [1] |
| *B. neotomae*-tdTomato | Transposon mutant of 5K33 having proD/tdTomato-nat genes | |
| *B. neotomae* ΔvirB4 | virB4 in-frame deletion mutant of B. neotomae 5K33 | |
| *Escherichia coli* NEB-5α | *fhuA2 Δ(argF-lacZ)U169 phoA glnV44 Φ80Δ(lacZ)M15 gyrA96 recA1 relA1 endA1 thi-1 hsdR17* | NEB |
| *E. coli* Ec100D pir-116 | *F- mcrA Δ(mrr-hsdRMS-mcrBC) ϕ80dlacZΔM15 ΔlacX74 recA1 endA1 araD139 Δ(ara, leu)7697 galU galK λ- rpsL nupG pir-116(DHRF)* | Bioresearch Technologies |
| *E. coli* β2155 | SM10 λpir derivative; *ΔdapA::erm* DAP auxotroph donor strain for conjugation | [2] |
| **Plasmids** | | |
| pMAR2xT7 | $Amp^R$, Himar1 transposase | [3] |
| pSR47s | R6K, sacB, $Km^R$, suicide vector | [4] |
| pBMTL2 | $Km^R$, broad host range vector | [5] |
| pBMTL3 | $Cam^R$, broad host range vector | [5] |
| pBAD | $Amp^R$, protein expression vector | Addgene (#37129) |
| **Cell Lines** | | |
| *Cell Lines* | *Relevant characteristics* | *Source* |
| J774A.1 | Mouse macrophage cell line, ATCC TIB-67 | ATCC |
| **PCR primers** | | |
| *Name* | *Sequences* | *Characteristics* |
| NatpMar2X-F | CCCGGTCTGGAGGACAGTAATGGGTACGACCCTTGATGACA | Amplification of NAT gene for pMAR2X |
| NatpMar2x-R | GTTGGCTGATAAGTCCCCGGTCTTTACGGGCAGGGCATGCTCATG | |
| pMAR2xNat-F | GGGGACTTATCAGCCAACCTGTTCCGACAGGGCCCAATTCGCCCTATAG | Amplification of pMAR2X for NAT gene |
| pMAR2xNat-R | TACTGTCCTCCAGACCGGGGACTTATCAGCCAACCTGTTCCGACA | |

| | | |
|---|---|---|
| trpD-1F | CGC *GGA TCC* AAG GTG AAC CAG GCC GTC | *trpD* in-frame mutant |
| trpD-1R | CCC ATC CAC TAA ACT TAA ACA ACC GGC GAT TTC CGG CAC | |
| trpD-2F | TGT TTA AGT TTA GTG GAT GGG CTG CTC AAT TCA GGC GCC | |
| trpD-2R | CGC *GAG CTC* AGG CTT CGA TCT TGC GAA | |
| trpDc-F | CGC *TCT AGA* ATGGCTGATTTGAAACCC | *trpD* complementation |
| trpDc-R | CGC *GAT ATC* TCAGGCCGGCTTGTCGTT | |
| metH-1F | CGC *GGA TCC* GGC ATT ACA ACC AGA CAA | *metH* in-frame mutant |
| metH-1R | CCC ATC CAC TAA ACT TAA ACA GGT CAA GGT CAG AAG ATC | |
| metH-2F | TGT TTA AGT TTA GTG GAT GGG GTG TCG GGT CTC TAT ATC | |
| metH-2R | CGC *GAG CTC* GCT GTT TTG CCG TTT CCT | |
| metHc-F | CGC *TCT AGA* ATGGCGTCTTCCCTTGAC | *metH* complementation |
| metHc-R | CGC *GAT ATC* TCAGGCCGCTTCGTCTTT | |
| qmetH-F | GCCAGCTTCAGGGCAATA | *metH* qPCR |
| qmetH-R | CGATGGAAGTGGAGGAGAAAG | |
| hisG-1F | CGC *GGA TCC* GACCATGCTTGGCGCTTG | *hisG* in-frame mutant |
| hisG-1R | CCC ATC CAC TAA ACT TAA ACA CCTCGCCTTCGACGCGCG | |
| hisG-2F | TGT TTA AGT TTA GTG GAT GGG ATCACCTCAACCGGCTCCAC | |
| hisG-2R | CGC *GAG CTC* GCTGAACCGTCGGCCACT | |
| hisGc-F | CGC *TCT AGA* ATGAGCGTCACATTGGCA | *hisG* complementation |
| hisGc-R | CGC *GAT ATC* TTATATTTCCAGTCCCGC | |
| qhisG-F | CGGATGACGACCGCAATTA | *hisG* qPCR |
| qhisG-R | AGATCGACGCTGCCATAAC | |
| hisD-1F | CGC *GGA TCC* CGTATGAGAACGCCCTCC | *hisD* in-frame mutant |
| hisD-1R | CCC ATC CAC TAA ACT TAA ACA CTCGCGGCGTACACGATCCA | |
| hisD-2F | TGT TTA AGT TTA GTG GAT GGG CGTTTCTCCTCCGGCCTT | |
| hisD-2R | CGC *GAG CTC* CGGGCATTGGGAACGTCG | |
| virB4q-F | GGCACTGAAACCATCGAGATAC | *virB4* qPCR |
| virB4q-R | GCTATACATCAGGCCGTTCAAA | |
| PvirB1-F | TGCCGCCTTGTTCACCGG | *virB1* promoter region |
| PvirB1-R | AGGATCGTCTCCTTCTCA | |
| rsh-1F | CGC *GGA TCC* CGGTTCGAGGGAGTTTCG | *rsh* in-frame mutant |
| rsh-1R | CCC ATC CAC TAA ACT TAA ACA GATTGTCGCCTCGTCCAA | |
| rsh-2F | TGT TTA AGT TTA GTG GAT GGG TTGGCGGAAATTGCCCAG | |
| rsh-2R | CGC *GAG CTC* AGGCCGGAGGTGAGGATA | |
| rshc-F | CGC *TCT AGA* ATGATGCGCCAATATGAG | *rsh* complementation |
| rshc-R | CGC *GAT ATC* CTATCCGTTCACACGCTT | |
| rshq-F | TGCGCGTTCTTCTGGTAAA | *rsh* qPCR |
| rshq-R | CATGGTCTCCTCGGCAATAC | |

| | | |
|---|---|---|
| deoR-1F | CGC *GGA TCC* GCA AAA CGG AAA GCG GCC | deoR/*glpR* in-frame mutant |
| deoR-1R | CCC ATC CAC TAA ACT TAA ACA CTG TTC TTC CAG CGA ATG | |
| deoR-2F | TGT TTA AGT TTA GTG GAT GGG GTC GTT ACC GAT GCG GGC | |
| deoR-2R | CGC *GAG CTC* GAT CAA ACG ACT GCC CAG | |
| lysR-1F | CGC *GGA TCC* ATTCATTCCATCGCATAA | *lysR*-family in-frame mutant |
| lysR-1R | CCC ATC CAC TAA ACT TAA ACA AAGATTGCGAATGGCGGA | |
| lysR-2F | TGT TTA AGT TTA GTG GAT GGG CGCCAGTTTCTCATGCTG | |
| lysR-2R | CGC *GAG CTC* GAATATAATTTATATTGTGG | |
| bvrR-1F | CGC*GGATCC*AGATTTGAACCGGTACAT | *bvrR* in-frame mutant |
| bvrR-1R | CCC ATC CAC TAA ACT TAA ACA CGGCATCTTGATATCGAA | |
| bvrR-2F | TGT TTA AGT TTA GTG GAT GGG GTGGATGATCGCACCATC | |
| bvrR-2R | CGC*GAGCTC*ATTGCTCAACGCGCCGTC | |
| PbvrR-1F | GGAAATCGAAGCGGCCTT | *bvrR* promoter region |
| PbvrR-2F | GGTGTGGAAAACCGCAAA | |
| PbvrR-R | GAGGTCAGGATGTTGCGG | |
| NHBvrR-F | CGC *AGATCT* AAGGAAGCTTCGGCAACG | *bvrR*_N'-His expression |
| NHBvrR-R | CGC *GAATTC* TTACGCTTCCCGGAAACG | |
| CHBvrR-F | CGC *CCATGG* ATGAAGGAAGCTTCGGCA | *bvrR*_C'-His expression |
| CHBvrR-R | CGC *GAATTC* TCAGTGGTGGTGGTGGTGGTGCGCTTCCCGGAAACGATAA | |
| pBAD33-F | GGAAGCGTAAGTCGACCTGCAGGCATGCAA | pBAD33 for *bvrR* conditional knockout |
| pBAD33-R | GGTTAATTCCTCCTGTTAGCTAGCCCAAAAAAACGGGT | |
| bvrR-F | TAACAGGAGGAATTAACCATGAAGGAAGCTTCGGCAAC | *bvrR* conditional knockout |
| bvrR-R | GCAGGTCGACTTACGCTTCCCGGAAACGAT | |
| bvrS-F | CGC *AGATCT* GTCGCAGAGACCCAGAAA | *bvrS* expression |
| bvrS-R | CGC *TTCGAA* TCATGCGCTGGCCGGAAG | |
| bvrRc-F | CGC *TCT AGA* ATGAAGGAAGCTTCGGCA | bvrR complementation |
| bvrRc-R | CGC *GAT ATC* TTACGCTTCCCGGAAACG | |
| ompR1-1F | CGC *GGA TCC* CAG CCA TAA TCT GGG ATG | *ompR1* in-frame mutant |
| ompR1-1R | CCC ATC CAC TAA ACT TAA ACA GCC AGC CAT CGT GAC GCG | |
| ompR1-2F | TGT TTA AGT TTA GTG GAT GGG CCC GCA AAC CCG GTC TGG | |
| ompR1-2R | CGC *GAG CTC* GCT TTC CAT GAA CCA CAT | |
| ompR1-F | CGC *AGATCT* GCTGCAGAAGAAGAAAAG | *ompR1* expression |
| ompR1-R | CGC *GAATTC* CTATTCGATGCTGAGTTT | |
| ompR1c-F | CGC *TCT AGA* ATGGCTGCAGAAGAAGAA | ompR1 complementation |
| ompR1c-R | CGC *GAT ATC* CTATTCGATGCTGAGTTT | |
| PompR1-F | CATGGTCAATCCCGAGCG | *ompR1* promoter |
| PompR1-R | CGAAACCCGCTATTGGTC | |
| PEnvZ1-F | CGC *AGATCT* AGATCGCCCTTCAGAAGA | |

| | | |
|---|---|---|
| PEnvZ1-R | CGC *GAATTC* TCAAGCCGGAATATGAAT | *envZ1* expression |
| rpoN-1F | CGC *GGA TCC* CCGGTTCCCACTTTTGGG | *rpoN* in-frame mutant |
| rpoN-1R | CCC ATC CAC TAA ACT TAA ACA CGTCATCTGTAGCAGCTTGA | |
| rpoN-2F | TGT TTA AGT TTA GTG GAT GGG GCCATCGTGGATGCGCTG | |
| rpoN-2R | CGC *GAG CTC* GGCTTGATGATTTCGGCT | |
| vjbR-1F | CGC *GGA TCC* GATCTCGTTCATTTTCCG | *vjbR* in-frame mutant |
| vjbR-1R | CCC ATC CAC TAA ACT TAA ACA CTGAAAAACCGGGTCAAT | |
| vjbR-2F | TGT TTA AGT TTA GTG GAT GGG GAAATCGCCGAAATCCTC | |
| vjbR-2R | CGC *GAG CTC* GAGCTTTTCTTTTCGCCT | |
| vjbR-F | CGC *AGATCT* GCGCTTCTAACCCGCATCCGG | *vjbR* expression |
| vjbR-R | CGC *GAATTC* TCAGACGAGATGCTGTACCTC | |
| PvjbR-F | CTTCGGTGCGCTTGCGGA | *vjbR* promoter region |
| PvjbR-R | AGTATCGCTTTGAAAGGA | |
| vjbRc-F | CGC *TCT AGA* ATGGCGCTTCTAACCCGC | vjbR complementation |
| vjbRc-R | CGC *GAT ATC* TCAGACGAGATGCTGTAC | |
| aqpZ-1F | CGC GGA TCC GAT TCG CAT ACT TGC CGT | aqpZ in-frame mutant |
| aqpZ-1R | CCC ATC CAC TAA ACT TAA ACA GGT GAG GAC GGT TAA ACC | |
| aqpZ-2F | TGT TTA AGT TTA GTG GAT GGG CGC TCG ACC GGC GTT GCC | |
| aqpZ-2R | CGC GAG CTC TAT TCG AAT CCG GCC CAT | |
| aqpZc-F | CGC *TCT AGA* ATGTTGAACAAATTATCG | aqpZ complementation |
| aqpZc-R | CGC *GAT ATC* TTAATCTCGGCCGAGCAG | |
| NLuc-F | CGC *TCTAGA* TAA GGA GGA AAA AAA ATGGTCTTCACACTCGAA | pBMTL3-NLuc |
| NLuc-R | CGC *AAGCTT* TTACGCCAGAATGCGTTC | |
| NLucPvirB1-F | CGC *GGATCC* TGCCGCCTTGTTCACCGG | pBMTL3-PvirB1_NLuc |
| NLucPvirB1-R | CGC *TCTAGA* AGGATCGTCTCCTTCTCA | |
| NLucPbvrR-F | CGC *GGATCC* GGAAATCGAAGCGGCCTT | pBMTL3-PbvrR_NLuc |
| NLucPbvrR-R | CGC *TCTAGA* GGTGTGGAAAACCGCAAA | |
| qrpoB-F | TCAGCGCGATCTGACTTATTC | *rpoB* qPCR |
| qrpoB-R | CTGCTCCTTGATGTCCTTGAT | |
| qompR-F | GCAGCCTTCTCTCACAATATCT | *ompR* qPCR |
| qompR-R | CGAGAATCAGAAGGTCGAAGTC | |
| qbvrR-F | TTATCGCGTCGAAACCTATACC | *bvrR* qPCR |
| qbvrR-R | ATGCGCGGCATCTTGATA | |
| qvjbR-F | AGCCGATCTGACTGTTCTTATG | *vjbR* qPCR |
| qvjbR-R | GTAATACGGAGCGTCTTCCTG | |

1. The underlined italicized sequence indicates the restriction enzyme recognition site.
